# Mutation of negative regulatory gene *CEHC1* encoding an FBXO3 protein results in normoxic expression of *HYDA* genes in *Chlamydomonas reinhardtii*

**DOI:** 10.1101/2024.03.22.586359

**Authors:** Xiaoqing Sun, Matthew LaVoie, Paul A. Lefebvre, Sean D. Gallaher, Anne G. Glaesener, Daniela Strenkert, Radhika Mehta, Sabeeha S. Merchant, Carolyn D. Silflow

## Abstract

Oxygen is known to prevent hydrogen production in Chlamydomonas, both by inhibiting the hydrogenase enzyme and by preventing the accumulation of HYDA-encoding transcripts. We developed a screen for mutants showing constitutive accumulation of *HYDA1* transcripts in the presence of oxygen. A reporter gene required for ciliary motility, placed under the control of the *HYDA1* promoter, conferred motility only in hypoxic conditions. By selecting for mutants able to swim even in the presence of oxygen we obtained strains that express the reporter gene constitutively. One mutant identified a gene encoding an F-box only protein 3 (FBXO3), known to participate in ubiquitylation and proteasomal degradation pathways in other eukaryotes. Transcriptome profiles revealed that the mutation, termed *cehc1-1*, leads to constitutive expression of *HYDA1* and other genes regulated by hypoxia, and of many genes known to be targets of CRR1, a transcription factor in the nutritional copper signaling pathway. *CRR1* was required for the constitutive expression of the *HYDA1* reporter gene in *cehc1-1* mutants. The CRR1 protein, which is normally degraded in Cu-supplemented cells, was stabilized in *cehc1-1* cells, supporting the conclusion that *CEHC1* acts to facilitate the degradation of CRR1. Our results reveal a novel negative regulator in the *CRR1* pathway and possibly other pathways leading to complex metabolic changes associated with response to hypoxia.

## Introduction

Using diverse physiological strategies, the unicellular photosynthetic alga *Chlamydomonas reinhardtii* acclimates to fluctuations in environmental factors including oxygen and carbon dioxide content; light quality and quantity; and concentrations of salt and organic molecules in the soil/freshwater habitats where it grows (reviewed by Johnson and Alric, 2013; Saroussi et al. 2017; Burlacot and Peltier, 2018; Burlacot et al. 2019). One acclimation strategy is to use hydrogenases to maintain redox balance in photosynthetic and fermentative metabolism, processes that have been investigated in detail using the powerful experimental tools of this system. Hydrogenases evolved as components of anaerobic energy metabolism in the evolution of early eukaryotes during the period of low atmospheric O_2_ and have been retained during evolution of most unicellular eukaryotic lineages including certain algae (Gould et al. 2019). Chlamydomonas cells accumulate O_2_-sensitive [Fe-Fe] hydrogenases in the chloroplast stroma where the enzymes transfer electrons from ferredoxin1 (PETF1, Fd) to protons, generating H_2_ (Winkler et al., 2010; Sawyer et al. 2017; Sawyer and Winkler, 2017).

Numerous studies have revealed the role of hydrogenases in pathways of hydrogen production (reviewed by Catalanotti et al. 2013; Grechanik and Tsygankov, 2022; King et al. 2022). Two pathways for hydrogen production utilize the photosynthetic electron transport chain. In direct biophotolysis, electrons from water splitting at PSII pass through the photosynthetic electron transport chain to PSI where light activation generates reduced Fd (Kosourov et al. 2020). In indirect biophotolysis, electrons from starch breakdown followed by glycolysis enter the electron transport chain at the level of plastoquinones via NAD(P)H dehydrogenase (NDA2) (Mus et al. 2005; Jans et al. 2008; Mignolet et al. 2012; Baltz et al. 2014) and are transferred from PSI to Fd. The production of hydrogen relieves the stress of excess reducing units in the stroma, thereby contributing to redox balance. Photoproduction of hydrogen occurs for short periods after initial illumination of cells following dark hypoxia, as would occur during the diurnal cycle (Gaffron and Rubin,1942; Ghirardi et al. 2007a). Hydrogenase reduction of protons via Fd provides an electron sink that stimulates ATP synthesis and photosynthesis during the dark to light transition. Ghysels et al. (2013) observed that the electron transport chain becomes re-oxidized and begins O_2_ evolution more rapidly in wild-type cells than in cells defective in hydrogenase. A third pathway for hydrogen production occurs in periods of dark hypoxia when fermentative metabolism is triggered (Grossman et al. 2010; Philipps et al. 2011; Atteia et al. 2013; Catalanotti et al. 2013; Yang et al. 2015). Fermentation utilizes carbon stores that accumulate during photosynthetic CO_2_ fixation in the light. Sugars derived from starch breakdown are utilized in glycolysis to produce ATP and pyruvate, the substrate for several fermentative pathways that regenerate NAD^+^ and produce acetate as well as toxic compounds including formate and ethanol. The enzyme pyruvate ferredoxin reductase (PFR1) also metabolizes pyruvate, yielding CO_2_ and reduced Fd. As in the photosynthetic pathways, Fd provides electrons to hydrogenases, which produce nontoxic H_2_ (van Lis et al. 2012; Noth et al. 2013).

Given the multiple roles for hydrogenases in photosynthetic and fermentative metabolism, it is likely that complex mechanisms have evolved for regulating expression of genes involved in hydrogen production. Dark hypoxia treatment induces increased expression of many genes as shown by analyses of transcript abundance (Mus et al. 2005; Hemschemeier et al. 2013; Subramanian et al. 2014; Blaby-Haas et al. 2016) and by quantitative proteomic analyses (Terashima et al. 2010; Nikolova et al. 2018). Among the genes whose expression is upregulated are nuclear genes *HYDA1* and *HYDA2*, encoding [Fe-Fe]-hydrogenases (Happe and Kaminski, 2002; Forestier et al. 2003) as well as *HYDG1* and *HYDEF1* genes, encoding assembly factors (Posewitz et al. 2004) needed for hydrogenase assembly and activity in the chloroplast stroma (Sawyer et al. 2017). *HYDA1* and *HYDA2* transcript levels decrease within minutes in the presence of oxygen. Reporter gene constructs have been used to demonstrate that the increase in *HYDA1* transcripts in response to hypoxia is regulated at the level of transcription (Stirnberg and Happe, 2004; Sun et al. 2013). Although *HYDA1/2* gene expression is inhibited by oxygen, data from RNA-seq experiments with cells grown under aerobic conditions shows that genes encoding hydrogenases and hydrogenase assembly factors are expressed in a diurnal manner, peaking within a few minutes of dark onset when fermentation may begin and again just prior to onset of light when photosynthesis begins (Zones et al. 2015; Gould et al. 2019; Strenkert et al. 2019). The transcription factor *CRR1*, identified by its role in regulation of the response to copper depletion (Quinn et al. 2000), plays a role in regulating the response to hypoxia (Quinn et al. 2002). Its SBP domain binds to GTAC-containing elements in the *HYDA1* promoter (Pape et al. 2012) and *crr1* mutants show decreased ability to activate expression of *HYDA1* in response to hypoxia (Pape et al. 2012; Hemschemeier et al. 2013).

Producing substantial quantities of hydrogen as an energy source from green algae will require a deeper understanding of all components of the pathway leading to hydrogen production (Ghirardi et al. 2007b: Hemschemeier et al. 2009; Torzillo et al. 2014; Burlacot and Peltier, 2018, King et al. 2022). To develop industrial production of hydrogen, it will be necessary to overcome the inhibitory effect of oxygen. Oxygen inhibits hydrogenase activity at both the post-transcriptional and transcriptional levels. The activity of [Fe-Fe] hydrogenase enzymes is acutely sensitive to oxygen. Under aerobic conditions in the light, O_2_ produced by water splitting at PSII leads to inactivation of the enzymes and cessation of H_2_ production. To utilize the hydrogen-producing capability of Chlamydomonas, it will be important to develop a better understanding of the regulatory mechanisms controlling expression of hydrogenase genes under different environmental conditions.

To identify components of the hypoxia signaling pathway(s) controlling hydrogenase expression, we designed a screen utilizing ciliary motility (Zhang and Lefebvre, 1997). Wild-type cells swim with two cilia that beat to propel the cells through liquid medium. Normal motility depends on the function of radial spokes, protein complexes that regulate dynein activity in the ciliary axoneme (Zhu et al. 2017). The *pf14* mutation results in ciliary paralysis due to a mutation in the *RSP3* gene encoding radial spoke protein 3 (Williams et al. 1989). Our screen used a reporter gene with a one kb fragment (including the 5’UTR) from upstream of the *HYDA1* gene to drive the expression of the *RSP3* gene (p*HYDA1-RSP3*; Sun et al. 2013). When expressed in *pf14* cells, the reporter gene conferred motility only in the absence of oxygen (Figure 1A). Transformant strains remained immotile under aerobic conditions but became motile after transfer to hypoxic conditions for 12 h, indicating that expression of the reporter gene was induced by hypoxia.

**Figure 1.**
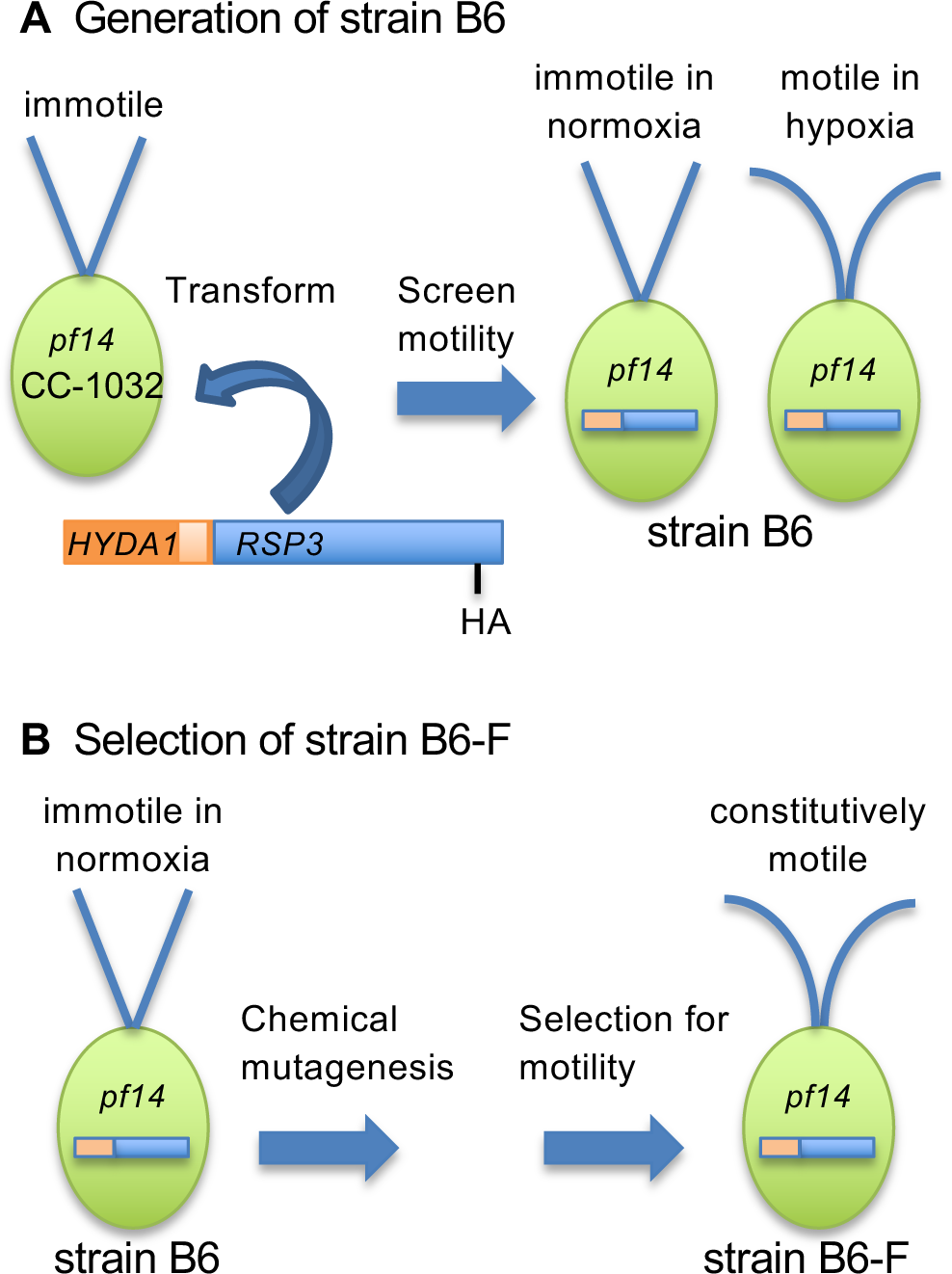
Generation of strains. **A)** Strain CC-1032 contains the *pf14* mutation in the *RSP3* gene, resulting in immotile (straight) cilia. It was transformed with a reporter gene construct in which a one-kb fragment containing the promoter (dark orange bar) and 5’ UTR (light orange bar) of the *HYDA1* gene was fused to the wild-type *RSP3* gene (blue bar) tagged with an HA-epitope. Transformed strain B6 (CC-4508) cells are immotile in aerobic conditions but become motile (curved cilia) in hypoxia due to expression of the *RSP3* reporter gene. **B)** After chemical mutagenesis, strain B6 cells were suspended in liquid media to select for cells able to swim to the top of the culture. One of these strains was designated strain B6-F (CC-4509).

Because the transformant strains could not swim in aerobic conditions, a simple and sensitive genetic screen was used to identify mutants defective in the suppression of *HYDA1* expression in oxygen. After spontaneous or chemical mutagenesis, the immotile cells were grown aerobically in liquid culture and screened for motile cells that could swim up to the meniscus (Fig. 1B; Sun et al. 2013). Many of these mutant cells had sequence alterations in the *HYDA1* promoter that allowed expression of RSP3 in the presence of oxygen. This screen also produced a constitutively motile mutant defective in a factor that acts in trans to regulate the activity of the *HYDA1* promoter in the transgene. The mutation, termed *cehc1* for <u>c</u>onstitutive <u>e</u>xpression of <u>h</u>ydrogenases and <u>c</u>opper-responsive genes, resides in a previously uncharacterized gene encoding an F-box protein in the *FBXO3* family (Kipreos and Pagano, 2000). We determined that this protein is a negative regulator of the *CRR1* transcription factor that controls expression of many genes in the copper assimilation pathway and the hypoxia response (Merchant et al. 2006; Sommer et al. 2010; Castruita et al. 2011; Pape et al. 2012; Hemschemeier et al. 2013). The *CEHC1* gene also appears to be involved in additional pathways controlling the response to hypoxia.

## Results

### Constitutively swimming mutant B6-F up-regulates expression of the *RSP3* reporter gene

Aerobically grown cells of the parental strain B6 (CC-4508; Supplemental Table 1), carrying one copy of the *pHYDA1-RSP3* reporter gene, showed the immotile phenotype expected of *pf14* mutants (Figure 1). After chemical mutagenesis, one mutant, B6-F, was motile and swam up to the meniscus from the bottom of the tube (Figure 2). Expression of the ectopic wild-type *RSP3* gene accompanied this motility, as shown by an immunoblot assay to detect the presence of the HA-tagged RSP3 protein (Figure 2B). The amount of the tagged protein in strain B6-F cells was increased compared with the parent B6 cells, consistent with the robust motility of the B6-F cells. The B6-F cells contained increased levels of the tagged protein in both cell bodies and cilia, where it was detected along the length and to the distal ends of the organelles (Figure 2C, arrowheads). The motility phenotype of strain B6-F correlates with and is likely dependent on the upregulated expression of the reporter gene.

**Figure 2.**
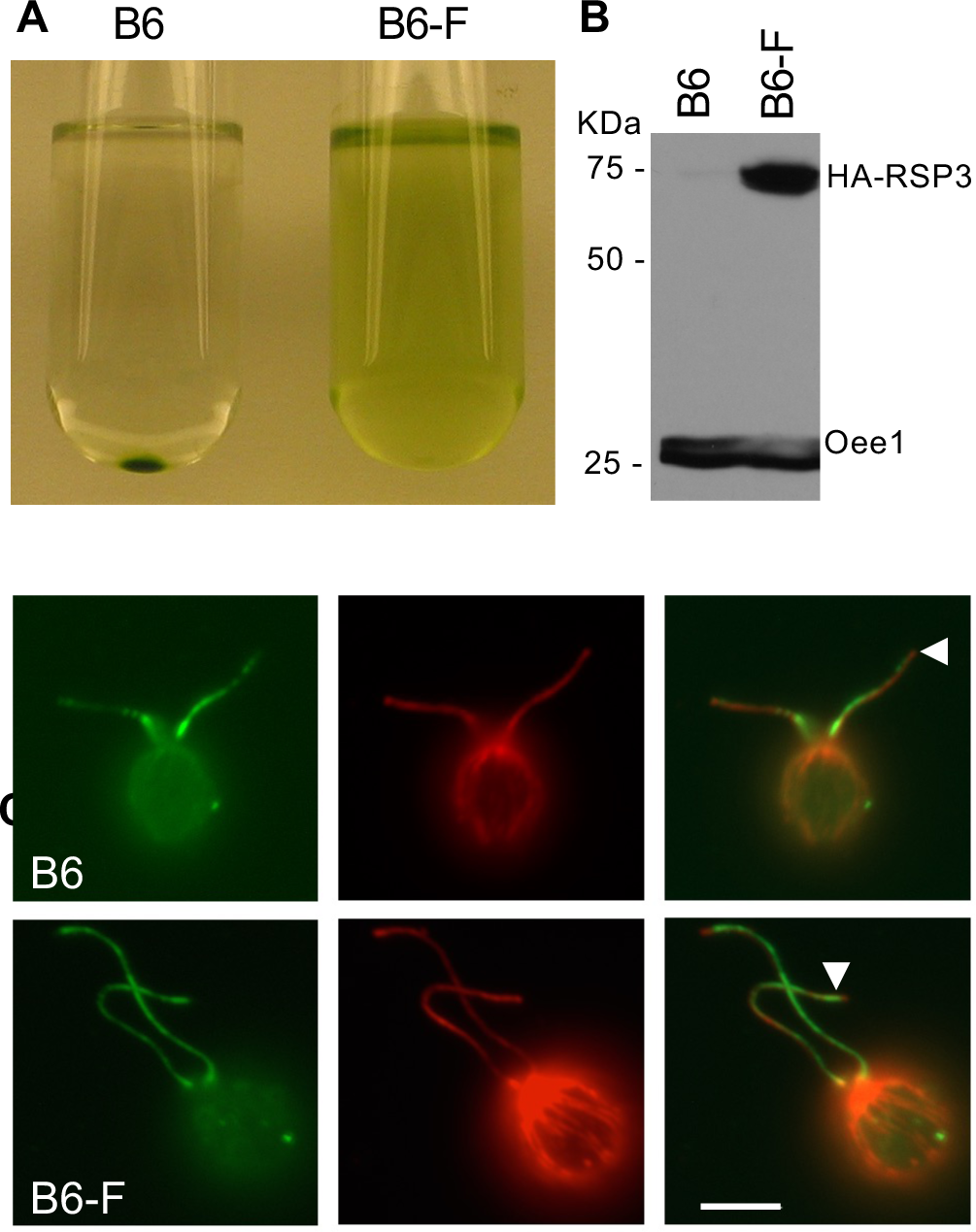
Constitutively swimming mutant strain B6-F expresses the *RSP3* reporter gene. **A)** Comparison of cell motility between strains B6 (CC-4508) and B6-F (CC-4509) in aerobic conditions. Equal numbers of cells from both strains were inoculated into liquid TAP medium in tubes with loose caps and kept under bright light for 5 days prior to photography. **B)** Immunoblot shows the relative level of HA-tagged RSP3 protein. Total protein from an equal number of aerobically grown cells was probed with anti-HA antibody using Immunoblot Analysis Method 1. Loading control was the Oee1 chloroplast protein. Positions of molecular weight markers are shown on the left. kDa, kiloDalton. **C)** Immunofluorescence images of cells from B6 and B6-F. The left panels show the presence of the RSP3 protein using an anti-HA primary antibody and a secondary antibody tagged with Alexa-Fluor 488 (green). Middle panels show cells stained with an anti-tubulin primary antibody and a secondary antibody tagged with Alexa Fluor 568 (red). Right panels show an overlay of the RSP3-HA and tubulin images. Scale bar represents 5 µm. Images are representative of >25 individual cells examined for each strain.

### A single mutation confers the motility phenotype on B6-F cells

A cross of motile B6-F (CC-4509) cells with immotile CC-613 cells, both of which contain an endogenous *pf14* mutation, showed that the new mutation in B6-F cells is linked within 25 cM of the reporter gene (Sun et al. 2013; see Methods). The *aphVIII* selectable marker gene conferring paromomycin resistance segregated 2:2 in all tetrads and segregated exclusively with motile cells in the PD (parental ditype) tetrads, indicating that the plasmid carrying the p*HYDA1-RSP3* reporter gene and the pSI103 plasmid (Sizova et al. 2001) used for co-transformation to generate strain B6, are tightly linked. An F1 progeny of the B6-F strain was back-crossed to the parent strain B6. The 2:2 segregation of motile to immotile progeny indicates that the causative mutation resided in a single genetic locus. For reasons described below, we designated the mutation as *cehc1* for <u>c</u>onstitutive <u>e</u>xpression of <u>h</u>ydrogenases and <u>c</u>opper responsive genes.

### The *cehc1* mutation maps to chromosome 1

We mapped the *cehc1-1* mutation using progeny from crosses to a polymorphic reference strain, CC-1952 and SNP markers (Figure 3). The mutation was mapped to a ∼1.1 Mb region of chromosome 1(Figure 3A, double-headed blue arrow; Supplementary Tables 2 and 3; Kathir et al. 2003). To identify the lesion within the mapped region, genomic DNA samples from strains B6 and B6-F were sequenced and the sequence was compared with the JGI v5.5 reference genome sequence (strain CC-503) (Merchant et al. 2007; Blaby et al. 2014). One SNP unique to B6-F within the 1.1 Mb region was detected, a T to C transition mutation localized to nucleotide 1,029,527 within gene Cre01.g005900 (Figure 3B). Consistent with the genetic linkage detected between the p*HYDA1-RSP3* reporter gene and the mutation leading to its constitutive expression, the site of insertion of the reporter gene construct was found on chromosome 1 at coordinate 5,034,215, or ∼4 Mb away (Figure 3A, black arrow).

**Figure 3.**
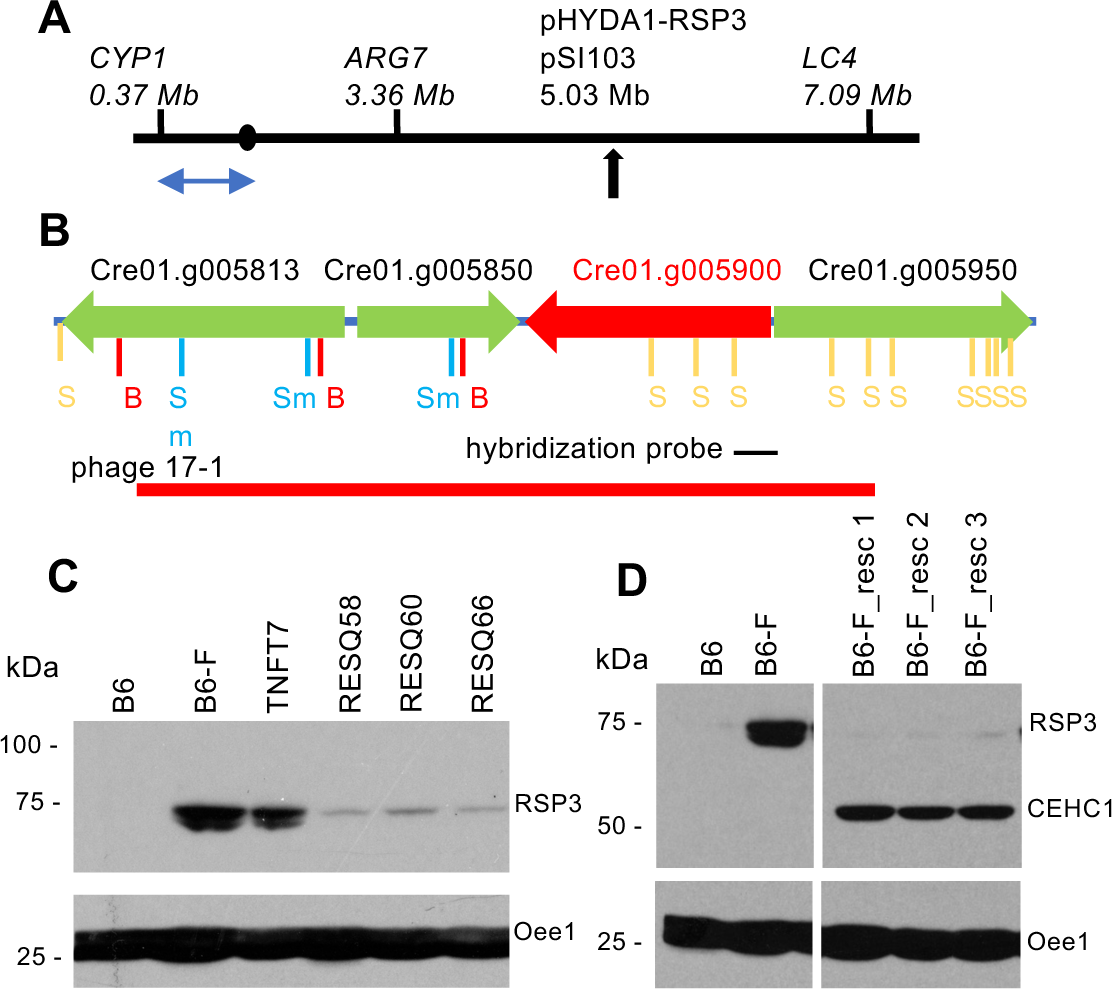
Mapping and phenotypic rescue of the *cehc1-1* mutation in strain B6-F. **A)** A partial map of chromosome 1 with the positions of three SNP mapping markers used with progeny from a cross of B6-F (CC-4509, laboratory strain background) x CC-2290 (polymorphic strain). Further mapping (Supplementary Table 2) localized the lesion in strain B6-F to the ∼1.1 Mb region indicated by the blue double sided arrow. A filled circle indicates the approximate location of the centromere; a black arrow indicates the site of insertion of the p*HYDA1-RSP3* reporter gene linked to the pSI103 selectable marker gene. **B)** Four gene models covering a 21.5 kb region of chromosome 1 (JGI ver 5.5 sequence), with the direction of the arrows representing the direction of transcription. The gene model in red contains the C to T mutation identified in the genome sequence of strain B6-F. Restriction sites are labeled beneath the gene models. (B, *Bam*HI; Sm, *Sma*I; S, *Sac*II). The 500 bp hybridization probe was used to screen a wild-type genomic DNA phage library to isolate phage 17-1. **C)** Protein from aerobically grown cells was processed for Immunoblot Analysis using Method 1. Immunoblot upper panel was probed with an anti-HA antibody to reveal the HA-tagged RSP3 protein encoded by the p*HYDA1-RSP3* reporter gene. Lower panel shows the loading control. Strains B6 and B6-F were controls. When transformed with an unrelated phage DNA the B6-F strain remained motile (TNFT7). When B6-F was transformed with phage 17-1 DNA pre-digested with *Bam*HI and *Sma*I, rescued (nonmotile) strains RESQ58/60/66 were obtained. Positions of molecular weight markers are shown on the left. kDa, kiloDalton. **D)** Immunoblot of total cell protein from three rescued (nonmotile) strains resulting from transformation of strain B6-F with a plasmid containing the cloned Cre01.g005900 gene tagged with the HA-epitope [pCEHC1-5.5H/S(1)3xHA5’]. Upper panel was probed with anti-HA antibody; lower panel, with anti-Oee1 antibody. The HA-tagged protein encoded by the Cre01.g005900 gene is labeled CEHC1.

### Gene Cre01.g005900 rescues the motility phenotype of *cehc1-1*

To determine whether the *cehc1-1* mutation is recessive, we generated stable diploid cells heterozygous for both *cehc1-1* and for the p*HYDA1/RSP3* reporter gene. These diploid strains showed the immotile phenotype, indicating that the reporter gene was not expressed and that the mutation is recessive (Supplementary Tables 4 and 5). We then used transformation to determine whether the SNP mutation in Cre01.g005900 is responsible for the motile phenotype of *cehc1-1* cells. A library of wild-type DNA cloned in a lambda phage vector was screened with a PCR-generated DNA fragment from the 3’ UTR of gene Cre01.g005900. Clone 17-1, which contained a 15.5 kb wild-type genomic fragment that covered the Cre01.g005900 gene model, the adjacent gene, and two partial gene models (Figure 3B) was transformed into B6-F cells along with the p*Hyg3* plasmid as a selectable marker (Berthold et al. 2002). Phenotypically rescued strains containing both mutant and wild-type copies of the *CEHC1* gene were expected to display the immotile *pf14* phenotype along with decreased expression of the HA-tagged *RSP3* reporter gene. Some transformant colonies had immotile cells (Supplementary Table 6) and some of these strains were tested for expression of the HA-tagged *RSP3* reporter gene using immunoblotting (Figure 3C). The HA-tagged RSP3 protein was expressed in strain B6-F and in a control strain transformed with an unrelated phage DNA (TFNT7). Expression of RSP3 was greatly reduced in three rescue strains (RESQ58, 60, 66) transformed with phage 17-1 DNA digested with BamH I/Sma I to disrupt the gene model to the left of Cre01.g005900 but to leave the Cre01.g005900 gene intact. Plasmid subclones containing only the Cre01.g005900 gene or an HA-tagged gene also resulted in transformants with a paralyzed phenotype (Supplementary Table 6). An immunoblot of proteins obtained from different rescue strains (B6-F_resc 1, 2, 3) expressing the HA-tagged copy of Cre01.g005900 showed that RSP3 protein expression was reduced compared to strain B6-F, whereas an HA-tagged protein of 55 kDa, the expected size for the protein encoded by Cre01.g005900, was expressed (Figure 3D). Thus, expression of the wild-type Cre01.g005900 gene prevents constitutive expression of the p*HYDA1-RSP3* reporter gene, leading to loss of motility and confirming that Cre01.g005900 is *CEHC1*.

### The *cehc1-1* lesion results in a splicing defect

Gene model Cre01.g005900 contains 13 introns, the first of which is affected by the T to C transition mutation in B6-F (Figure 4). The 5’ consensus donor splice site of the first intron is changed from GT to GC (Figure 4A). To determine whether the mutation affects splicing of the transcript, we used reverse transcription PCR to amplify a fragment spanning the intron (Supplementary Table 7). The resulting cDNA fragment from RNA of strain B6-F (*cehc1-1*) was larger than the fragment from strain B6 (Figure 4B). Sequencing of the cDNA verified that the intron 1 sequence of 86 bp was retained in transcripts from strain B6-F but was missing in transcripts from strain B6. The unspliced transcripts likely do not produce a functional product because an in-frame premature stop codon within the intron follows the SNP mutation. Re-initiation of translation at the first start codon downstream of the mutation (22 nt downstream, within intron 1) would produce a protein in the incorrect reading frame. Data from RNA-seq (see below) revealed that the mutant sequence can function at a low level as a splice donor, perhaps due to inherent flexibility of the splicing machinery (Papasaujas and Valcárcel, 2016). Excision of intron 1 was found in all transcripts in the wild-type strain (Figure 4C). In contrast, correct intron excision occurred for approximately 9% of transcripts in the mutant strain, showing that the *cehc1-1* mutation is a hypomorphic as opposed to a null allele. Overall transcript levels (spliced and unspliced) were two-fold higher in the mutant strain for reasons that are unclear.

**Figure 4.**
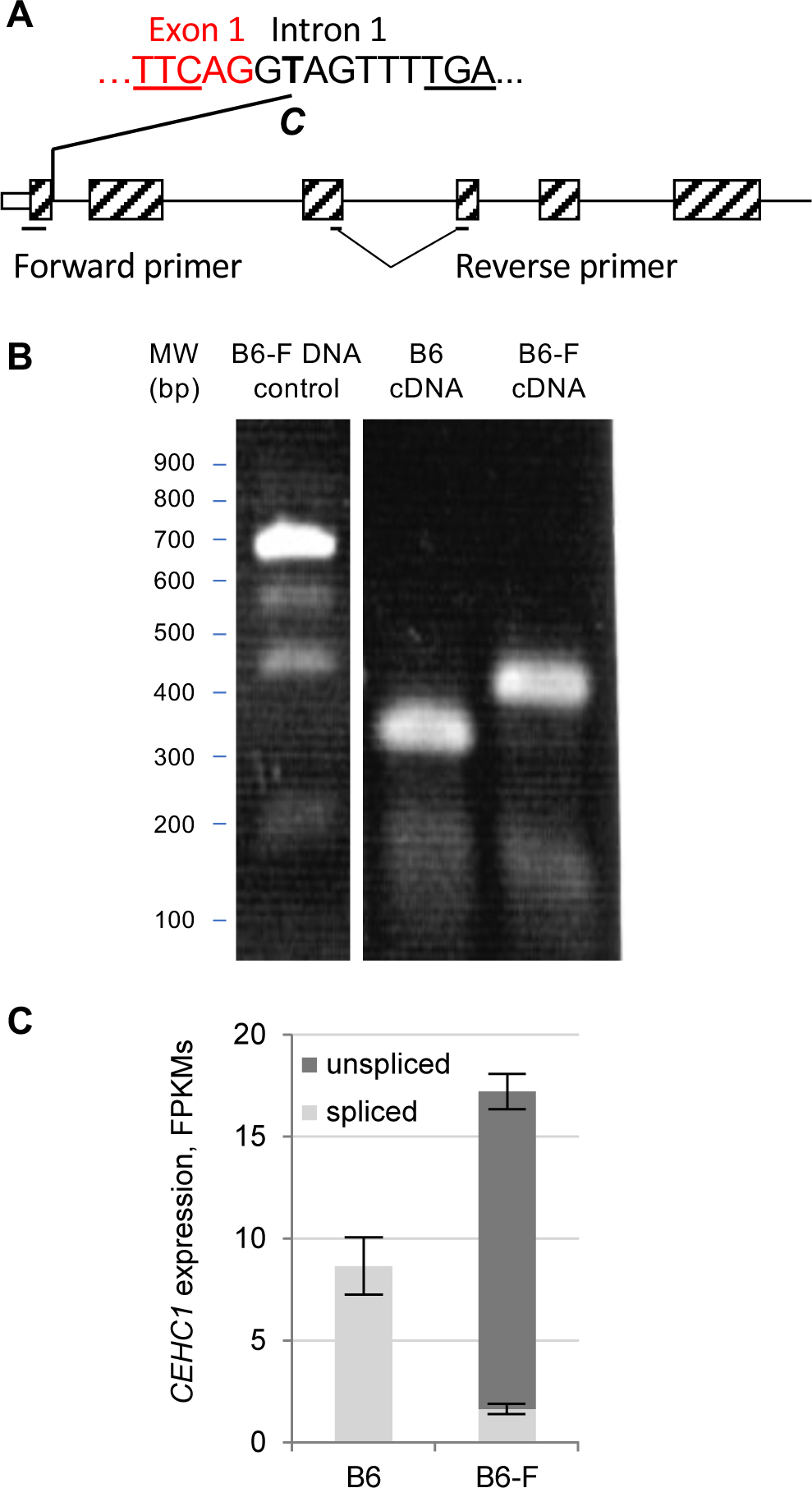
The *cehc1-1* mutation in strain B6-F disrupts splicing. **A)** Gene model diagram showing the position of the point mutation in the *CEHC1* gene (Cre01.g005900). The 5’UTR (blank rectangle), the first six exons (hatched rectangles) and the first five introns (solid lines) are depicted. The position of the T-C single nucleotide mutation at the second nucleotide from the 5’ end of intron 1 is indicated above the gene model (red: 3’ end of exon 1; black: intron 1). The red underline shows the reading frame; the black underline of TGA shows the in-frame stop codon. Forward and reverse primers used for reverse transcription PCR in panel B are marked (Supplementary Table 7). The reverse primer spans a splice site. **B)** Gel electrophoresis of RT-PCR products. Total RNA from aerobically grown B6 (CC-4508; *CEHC1*) and B6-F (CC-4509; *cehc1-1*) strains was reverse transcribed using the reverse primer as the gene specific primer and PCR amplified using the forward and reverse primers. The cDNA templates from strain B6 yielded a product smaller than the one from B6-F. Genomic DNA from strain B6-F was used as a negative control. **C)** RNA-Seq was used to quantify transcripts from the *CEHC1* gene from cultures of B6 and B6-F. The fraction of transcripts with and without intron 1 (unspliced versus spliced) was calculated from aligned fragments at the intron 1 splice donor site. Shown here is the mean ± standard deviation (*N* = 3) of transcript abundance in terms of fragments per kb per million mapped fragments (FPKMs).

### The *cehc1-1* mutation affects an FBXO3-like protein

Gene model Cre01.g005900 encodes a predicted protein of 474 amino acids with a molecular mass of 52,195 D and a pI of 6.34. The TargetP (Emanuelsson et al. 2000) and PSORT (http://www.psort.org/) (Horton et al. 2007) algorithms did not detect localization signals and predicted a cytosolic localization. Basic Local Alignment Search Tool (BLAST) searches showed that the amino acid sequence is conserved across a wide range of eukaryotes including metazoans, with the highest sequence similarity (35% – 45% identity) in proteins from the clade Viridiplantae including green algae and nonvascular and vascular land plants (Supplementary Figure 1). Analysis of the Chlamydomonas protein along with putative homologs from Arabidopsis and humans using InterPro 85.0 software (Blum et al. 2020) and Alphafold software (Varadi et al. 2022) identified three conserved domains in the same order (Figure 5). From N-terminus to C-terminus, these include an F-box or F-box-like domain, a KNR4/Smi1-like domain, and an ApaG domain. The *Homo sapiens* FBXO3 protein has been shown to be a component of an SCF type (Skp1, Cullin, F-box protein) ubiquitin E3 ligase (Ilyin et al. 2000; Chen et al. 2013). The Arabidopsis SKIP16 (SKP/ASK-interacting protein 16) was identified in a yeast-two hybrid screen to interact with ASK2, a Skp1 homolog (Risseeuw et al. 2003). These comparisons suggested that the Chlamydomonas CEHC1 protein may function in an SCF complex. A BLAST search of the Chlamydomonas genome showed that Cre01.g005900 is the only gene encoding a protein with all three domains.

**Figure 5.**
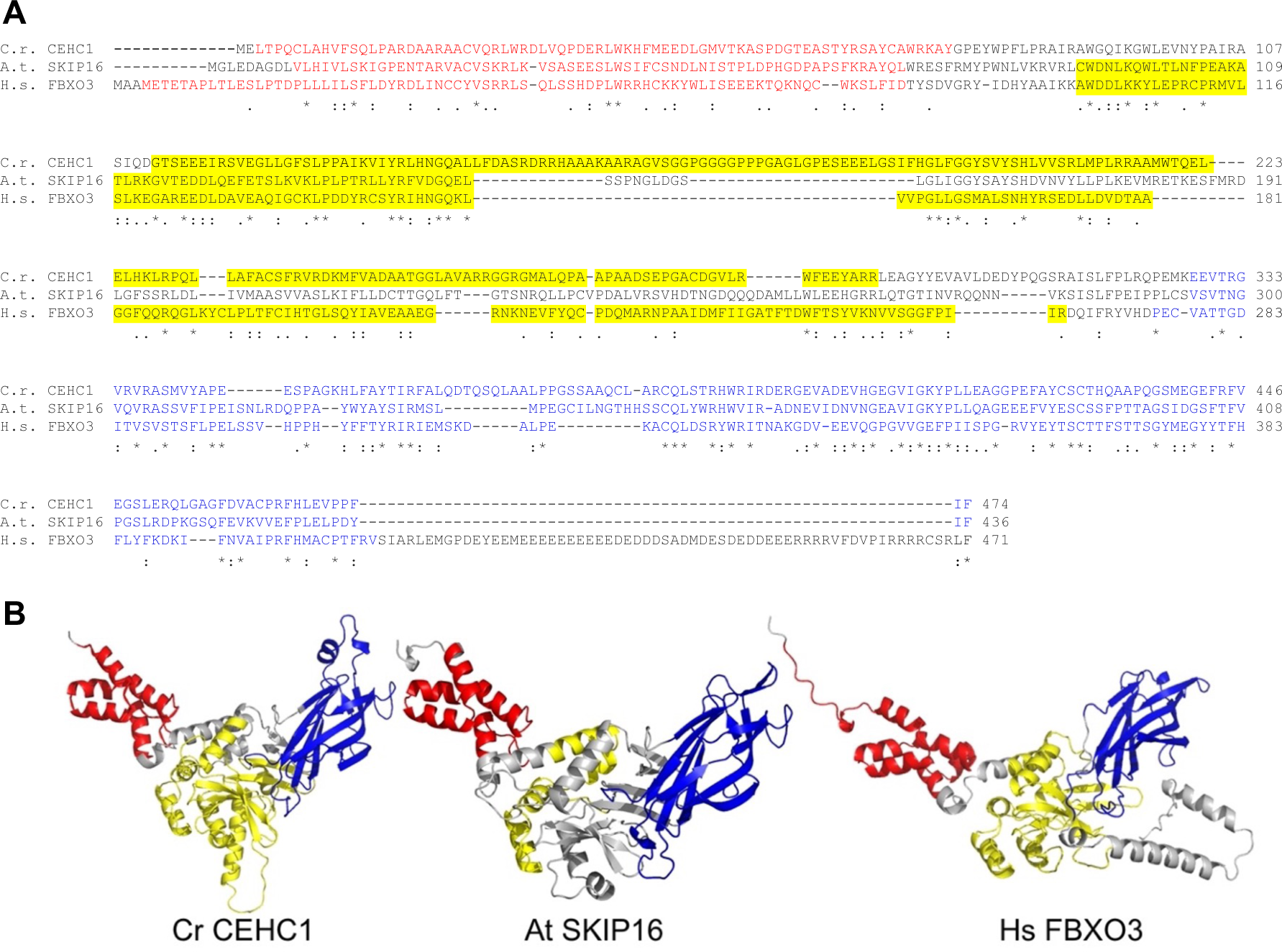
Conservation of FBXO3-like proteins **A)** The CEHC1 protein (*C. reinhardtii* locus Cre01.g005900) aligned with the *A. thaliana* SKIP16 protein (NP_563759.1) and the *H. sapiens* FBXO3 protein isoform 1 (NP_036307.2). Multiple sequence alignment utilized MUSCLE v3.8.31 software (EMBL-EBI). Conserved domains for each protein were analyzed using InterProScan v5.64-96.0 (EMBL-EBI). The F-Box/F-box-like domain is indicated in red text highlight, the KNR4/Smi1-like domain in yellow, and the ApaG domain in blue. **B)** AlphaFold structure predictions from UniProt (accessions: Cr CEHC1=A0A2K3E540, At SKIP16=Q9LND7, Hs FBOX3=Q9UK99) were visualized and conserved domains were colored as in panel A with PyMol (v2.5.0).

### The *cehc1-1* mutation alters gene expression

In *cehc1-1* mutant cells, the *RSP3* reporter gene is expressed constitutively from its activated *HYDA1* promoter. To determine whether the *cehc1-1* mutation results in changes in the expression of other genes, strains B6 (*CEHC1*), B6-F (*cehc1-1*), and B6-F_resc (*cehc1-1*; *CEHC1*-TG) (trans gene) were grown in triplicate under aerobic conditions and RNA was isolated using conditions to minimize exposure to hypoxia prior to cell lysis (Figure 6). Transcript levels for specific genes were assayed using qPCR (Figure 6A). All three strains showed similar levels of transcripts from the *AMYB1* gene encoding beta amylase, used as a control for detecting the intracellular O_2_ availability in sampled cells (Mus et al. 2007). Levels of *HYDA1* transcripts were elevated by ∼15-fold in *cehc1-1* cells relative to *CEHC1*. The *cehc1-1* mutation appears to be responsible for upregulated expression of *HYDA1* because rescue with the wild-type gene in strain B6-F_resc restored *HYDA1* transcripts to the low levels observed in *CEHC1* cells.

**Figure 6.**
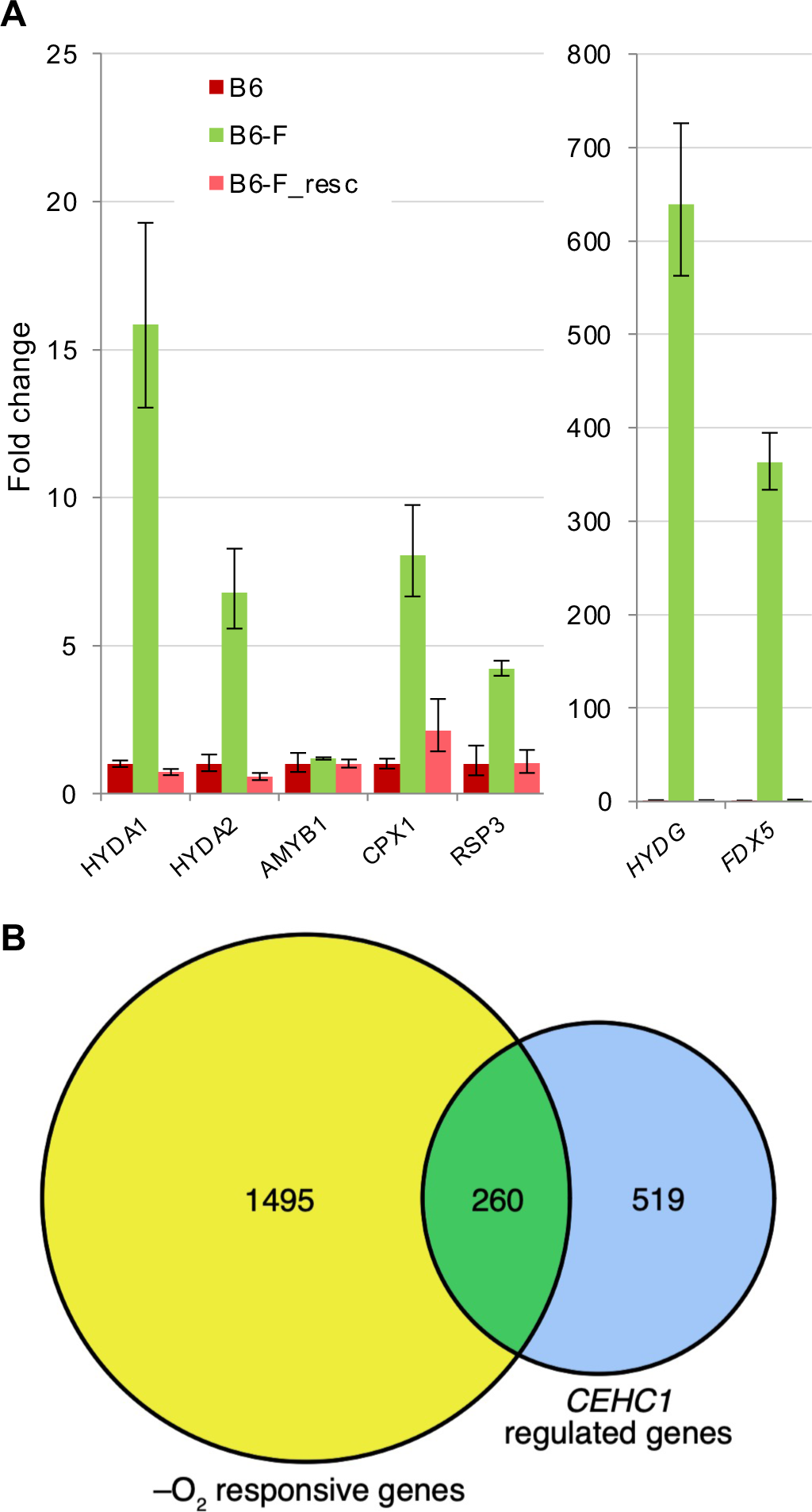
Analysis of transcript levels altered by *cehc1-1* mutation. RNA was isolated from triplicate cultures of strains B6 (*CEHC1;* CC-4507), B6-F (*cehc1-1*; CC-4509), and B-F_resc strain (*CC-5386*), grown aerobically in TAP media. **A)** Transcript levels were determined using qPCR assays (see Methods). Each mean transcript level was normalized to the house-keeping gene RCK1 (Cre06.g278222) and calibrated to the level in strain B6 set at 1. Genes indicated on the X axis encode: *HYDA1* hydrogenase Cre03.g199800; *HYDA*2 hydrogenase Cre09.g396600; *AMYB1*, beta-amylase Cre06.g307150; *CPX1*, coproporphyrinogen III oxidase Cre02.g085450; RSP3, radial spoke protein 3 Cre06.g291700; *HYDG1*, hydrogenase assembly factor Cre06.g296700; *FDX5*, apoferredoxin Cre17.g700950. **B)** RNA-Seq analysis of transcript levels altered by *cehc1-1* mutation compared with transcripts altered by dark hypoxia. RNA-Seq was performed on the same RNA samples to identify differentially expressed genes (DEGs) affected by *CEHC1* (see Methods). Genes upregulated or down-regulated in the *cehc1-1* mutant by at least two-fold (*CEHC1* regulated genes) were compared with genes upregulated or down-regulated by at least two-fold in a previous study conducted with a *CEHC1 (CRR1* rescued) strain exposed to aerobic and dark anoxic conditions (+/–O_2_). Transcript levels were reanalyzed using the latest JGI *C. reinhardtii* assembly (CC-4532 v6.1) and annotations to facilitate comparison with this study. The overlap in identified DEGs between the studies is presented here as a Venn diagram. See Supplementary Dataset 3.

Because the *HYDA1* promoter is known to be a target for upregulation by the *CRR1* transcription factor (Quinn et al. 2002; Pape et al. 2012), we examined expression of known *CRR1* target genes *CPX1* and *FDX5* (Kropat et al. 2005; Lambertz et al. 2010; Castruita et al. 2011; Pape et al. 2012; Kropat et al. 2015; Blaby-Haas et al. 2016). An increase in transcripts was observed in *cehc1-1* mutant cells; transcripts returned to low levels in the B6-F_resc strain. We also tested transcript levels for genes (or proteins) previously shown to be upregulated in hypoxia including *HYDA2* and *HYDG1* (Mus et al. 2007; Terashima et al. 2010; Hemschemeier et al. 2013; Blaby-Haas et al. 2016). Although not previously noted as targets of the *CRR1* transcription factor, both genes showed increased transcript levels in *cehc1-1* cells as compared to *CEHC1* and these levels were rescued by the wild-type gene.

Based on both motility and the production of the HA-tagged protein observed in *cehc1-1* cells (Figures 2 and 3), we expected that, of the two copies of the *RSP3* gene present in all three strains, the *RSP3* reporter gene under control of the *HYDA1* promoter would be upregulated only in the *cehc1-1* strain. The endogenous *RSP3* gene (Williams et al. 1989) produces a transcript characteristic of the wild-type gene, expected to be uniform among the *cehc1-1*, *CEHC1*, and rescue strains. The qPCR results showed that the *RSP3* transcript level in motile *cehc1-1* cells is elevated ∼4-fold compared to that of immotile *CEHC1* and rescue cells (Figure 6A). This relatively small change in transcript level associated with presence or absence of motility underscores the sensitivity of the *RSP3* reporter gene and the power of the motility assay for genetic discovery of negative regulators. Because the *cehc1-1* mutation leads to constitutively elevated expression of the *HYDA1* gene and other genes known to be targets of the transcription factor *CRR1*, a key regulator of the copper response pathway in Chlamydomonas, we designated the mutation as *cehc1* for <u>c</u>onstitutive <u>e</u>xpression of <u>h</u>ydrogenases and <u>c</u>opper responsive genes. Because the screen we employed is most useful for identifying negative regulatory genes, the mutants would be expected to show elevated expression of genes downstream of the negative element.

To gain a comprehensive view of the effect of the *cehc1-1* mutation on the transcriptome, the RNA samples were analyzed using RNA-seq and mapped to the JGI v6.1 genome (see Methods; Supplementary Dataset 1). The transcriptome data were subjected to three filters: genes whose expression differed by at least two-fold between the B6 (*CEHC1*) and B6-F (*cehc1-1*); genes for which the FPKMs were greater than 5.0 in at least one strain; genes whose altered expression in the *cehc1-1* mutant strain were rescued by at least 50% in the B6-F_resc strain. Transcripts from 279 genes upregulated in the *cehc1-1* mutant and 500 genes down-regulated in the mutant (Supplementary Dataset 2) met these criteria. The greatest differences in transcript levels observed between the *CEHC1* and *cehc1-1* strains were in the set of genes upregulated in the mutant; 89 genes were upregulated between 4- and >1×10^4^-fold in the mutant strain, whereas 37 were down-regulated, between 4- and 42-fold. Controls for the experiment included transcript levels for genes previously shown to have stable expression levels (Shi et al. 2017; Zones et al. 2015) (Supplementary Table 10). For six such genes, the *cehc1-1*/*CEHC1* ratio ranged from 1.5 to 0.7-fold. As noted above for qPCR results, the *AMYB1* transcript levels were consistent with exposure of all the cultures to similar levels of aeration. Also confirming the qPCR results is the relatively small increase in *RSP3* transcripts (∼2-fold) required to generate motility in the B6-F strain. Transcript levels for the *CEHC1* gene differed by 2-fold between the mutant and wild-type strains but were higher in the rescue strain as might be expected for the strain carrying an extra copy of the gene.

### Overlap with genes responding to dark hypoxia

The *cehc1-1* mutation alters expression of the hypoxia-induced *HYDA1* gene. We took advantage of a previous RNA-seq study by Hemscheimeier et al. (2013a) to identify other genes regulated by hypoxia that show altered expression in the *cehc1-1* mutant. Of the 779 genes regulated by at least 2-fold in aerobically grown *cehc1-1* cells, 260 genes overlapped with a set of genes showing at least 2-fold altered expression in the previous study of wild-type cells exposed to dark hypoxia (Figure 6B; Supplementary Dataset 3, columns F and V). The overlapping genes included 94 genes upregulated in the *cehc1-1* mutant and 166 down-regulated genes. The results confirm that CEHC1 plays a role in regulating expression of many but not all genes regulated by oxygen levels. The list of transcripts upregulated both by hypoxia and by the *cehc1-1* mutation includes a second hydrogenase gene, *HYDA2,* as well as *HYDG1* and *HYDEF1*, encoding hydrogenase assembly factors (Posewitz et al. 2004). Transcripts from three members of the HCP (hybrid cluster protein) gene family *(HCP1, HCP3 and, HCP4*), were upregulated in the *cehc1-1* mutant and returned to wild-type levels in the rescue strain. The HCP enzymatic activity includes conversion of nitric oxide into nitrous oxide and the enzyme may play a role in NO based signaling (Hagen, 2019). Increased expression of *HCP3* and *HCP4* in response to dark hypoxia was noted previously (Mus et al. 2007; Hemschemeier et al. 2013; Olson and Carter, 2016).

We found that the majority (> 90%) of the 166 genes that were downregulated by two-fold or more in the *cehc1-1* mutant were also repressed in dark hypoxia while only 13 genes showed upregulation in these conditions (Supplementary Dataset 3). These results imply that CEHC1 may function as a positive and a negative regulator at the same time, depending on the regulatory pathway.

### Overlap with CRR1 target genes

The *RSP3* reporter gene contains within its *HYDA1* promoter a CRR1 binding site or Cu responsive element (CuRE) (Kropat et al. 2005). We compared our RNA-seq data with published RNA-seq data identifying CRR1 target genes by comparing expression levels in a *crr1-2* mutant strain and a *CRR1*-rescued strain grown in media without copper (Castruita et al. 2011; Kropat et al. 2015; Blaby-Haas et al. 2016). A Venn diagram in Figure 7 (Supplementary Dataset 4) shows 126 genes upregulated or down-regulated by more than four-fold in the *cehc1-1* mutant strain (CEHC1-regulated genes). A subset of these genes with assigned symbols in the JGI v6.1 genome annotations are indicated in colored boxes. We observed overlap with 28 genes identified as CRR1 targets. Many of these genes have putative CuREs upstream of the genes. The RNA samples in our study were from cells grown in aerated, copper-replete medium, conditions that should suppress CRR1 activity. The finding that genes regulated by CRR1 are similarly regulated by loss of CEHC1 activity is consistent with the hypothesis that CEHC1 acts in the same pathway as CRR1 and that loss of CEHC1 function leads to increased CRR1 activity. The increased activity is likely due to a post-transcriptional mechanism because the levels of *CRR1* transcripts were relatively constant in the *CEHC1*, *cehc1-1*, and rescue strains (Supplementary Table 10). Relevant to this conclusion is the observation of little change in *CRR1* transcript levels in plus or minus copper, indicating that the copper response is regulated at the level of the CRR1 polypeptide (Kropat et al. 2005).

**Figure 7.**
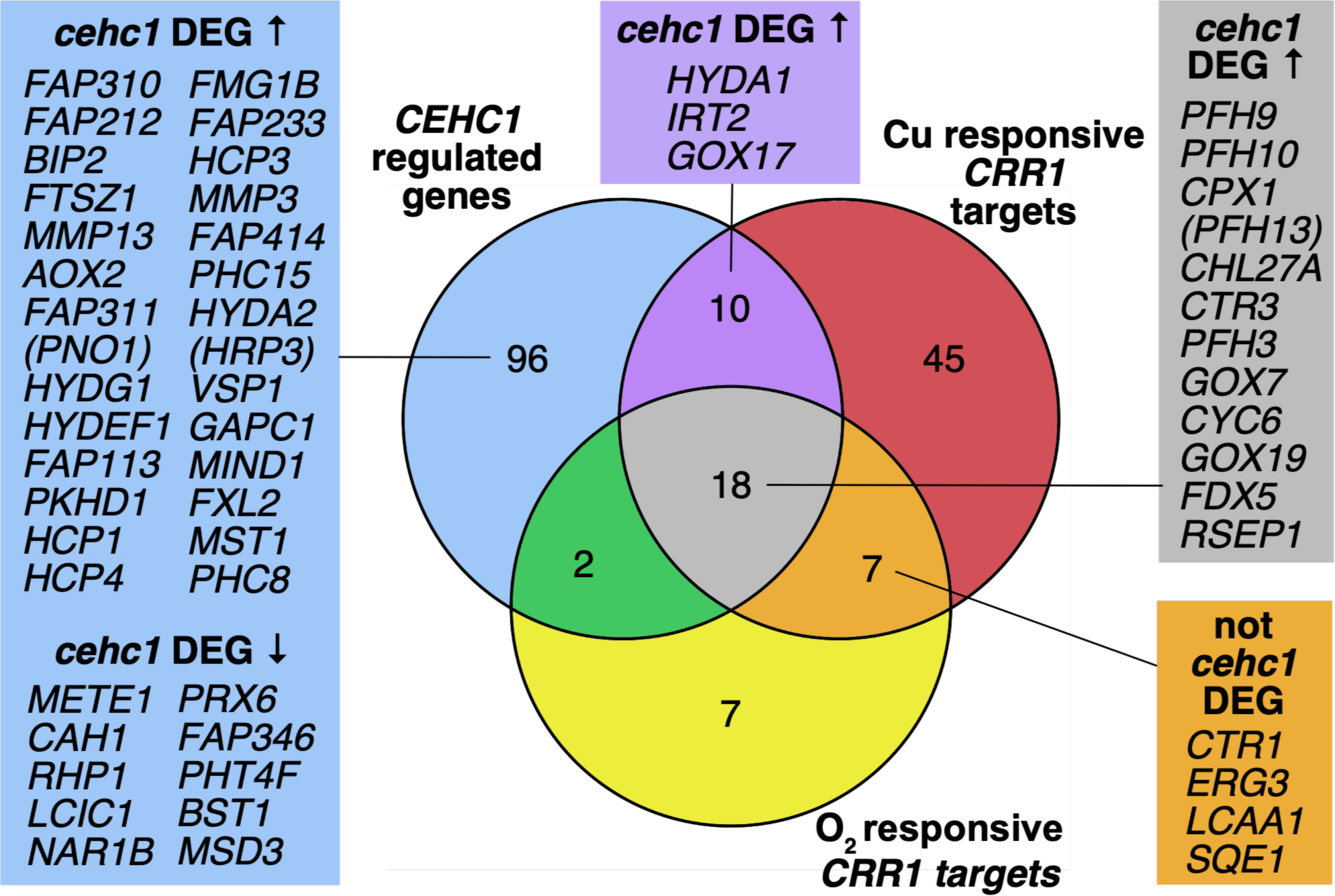
Overlap of differentially expressed genes in *cehc1-1* and *crr1-1* mutant strains. RNA-Seq was performed on B6 (*CEHC1*), B6-F (*cehc1-1*), and B6-F_resc (*CEHC1* rescued) strains in order to identify differentially expressed genes (DEGs) affected by *CEHC1* (see Methods). Genes upregulated or down-regulated in the mutant by at least four-fold and have assigned symbols in the JGI listing are shown. Two previous studies conducted on *crr1-2* mutant and *CRR1* rescued strains grown with and without Cu (+/–Cu) and under aerobic and dark anoxic conditions (+/–O_2_) were reanalyzed using the latest *C. reinhardtii* assembly and annotations to facilitate comparison with this study. The overlap in identified DEGs between the three studies is presented here as a Venn diagram. Genes with assigned gene symbols in several groups are highlighted, with deprecated gene symbols in parentheses. See Supplementary Dataset 4.

Many genes defined as CRR1 targets are also upregulated in response to dark hypoxia conditions (Quinn et al. 2002; Hemschemeier et al. 2013). Mutant *crr1* cells are unable to grow in oxygen-deficient medium, with or without copper (Eriksson et al., 2004). To assess the possible role of CRR1 in regulating genes in the CEHC1 pathway, transcript levels from the remapped Hemschemeier study (2013) in aerobic conditions vs dark anaerobic conditions were compared between a *crr1-2* mutant strain and a strain rescued with a wild-type *CRR1* copy. An O_2_ responsive CRR1 target was defined as a gene whose expression change in the *crr1-2* mutant was less than 25% of the expression change in the wild-type cells in the presence vs. absence of O_2_ (Fig 7; Supplementary Dataset 4, column AA). Eighteen O_2_-responsive CRR1 target genes overlapped with Cu-responsive CRR1 targets whose expression is altered by the *cehc1-1* mutation, reaffirming a role for CEHC1 in the CRR1 pathway.

### Interaction of the *CEHC1* and *CRR1* genes

To examine whether the *CEHC1* and *CRR1* genes function in the same signaling pathway, we determined the role of these genes in expression of the reporter and other genes. Strains were constructed with all four possible combinations of the wild-type and mutant alleles (Supplementary Tables 1, 5). Immunoblots of total cell proteins from aerobically grown cells revealed that expression of the reporter gene was elevated in strains with the *cehc1-1*; *CRR1* combination, the same combination that exists in the B6-F strain (Figure 8). Constitutive *RSP3* expression is lost in the presence of the *crr1-1* mutation (*cehc1-1*; *crr1-1*), indicating that *crr1-1* is epistatic to *cehc1-1*. That is, the phenotype of the *cehc1-1* mutant requires the action of the *CRR1* gene product. A similar epistasis was observed for the expression of endogenous genes upregulated in the *cehc1-1* mutant (Figure 8B). Progeny from a single tetratype tetrad were grown aerobically in copper-replete medium and RNA was isolated for qPCR analysis of relative transcript levels. Each strain expressed similar levels of *AMYB1* transcripts, indicating similar levels of aeration. As observed for expression of the RSP3 reporter protein, only the *cehc1-1*; *CRR1* strain showed upregulated expression of the *CRR1* target genes *HYDA1*, *FDX5* and *CPX1*, as well as of the hypoxia regulated genes *HYDA2* and *HYDG1*. The *crr1-1* mutation suppressed the *cehc1-1* mutant phenotype in double mutant progeny. The results confirmed that the up-regulation of the transcript levels of these endogenous genes in *cehc1-1* strains is dependent on *CRR1*.

**Figure 8.**
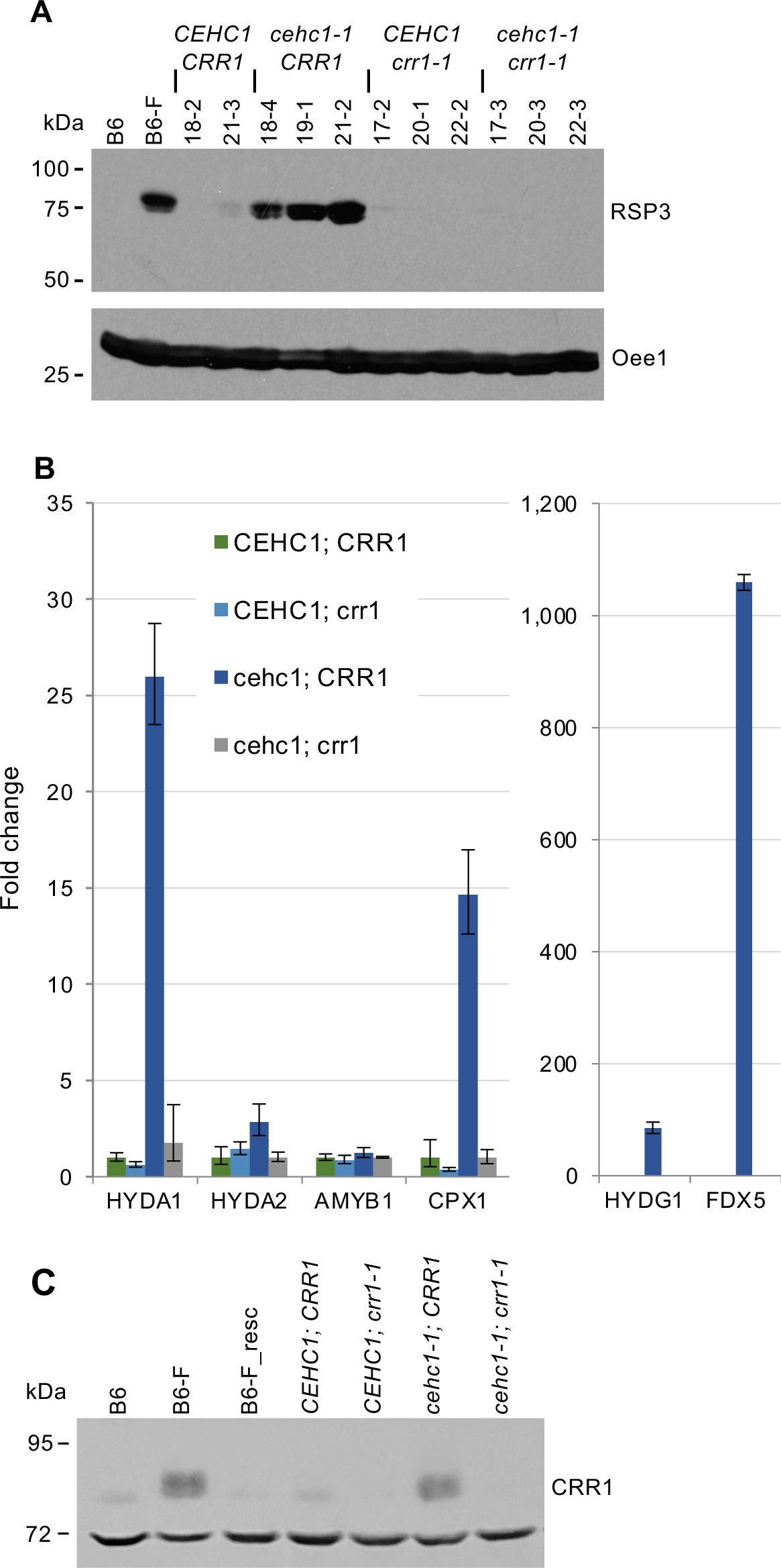
Interaction of *CEHC1* and *CRR1* genes. **A)** Immunoblot to examine RSP3-HA expression. Strain CC-4509 (*cehc1-1*) was crossed with CC-3959 (*crr1-1*). Progeny strains (18-2 to 22-3), all containing the *RSP3* reporter gene, were examined for alleles at *CEHC1* and *CRR1* (see Methods). Cultures were grown in aerated TAP media. Total protein from an equal number of cells was processed for Immunoblot Analysis using Method 1. The blot was probed with anti-HA antibody to detect the RSP3-HA protein (upper panel) and with an antibody against Oee1 (lower panel) as a loading control. Controls are strain B6 strain (CC-4507, *CEHC1; CRR1)* and strain B6-F (CC-4509, *cehc1-1; CRR1).* Genotypes of progeny strains are indicated (“XS” strains, Supplementary Table 1). **B)** q-PCR analysis to examine transcript levels in strains from one tetrad with different *CEHC1* and *CRR1* allele combinations (Supplementary Table 1). RNA was isolated from triplicate cultures of strains grown aerobically in TAP media. Levels of transcripts were determined in triplicate qPCR assays for each RNA sample (see Methods). Each mean transcript level was normalized to that of the house-keeping gene *RCK1* (Cre06.g278222), and calibrated to the level in the *CEHC1; CRR1* strain that was set at 1. *HYDA1* hydrogenase Cre03.g199800; *HYDA*2 hydrogenase Cre09.g396600; *AMYB1*, beta-amylase Cre06.g307150; *CPX1*, coproporphyrinogen III oxidase; Cre06.g291700; *HYDG1*, hydrogenase assembly factor Cre06.g296700; *FDX5*, apoferredoxin Cre17.g700950. **C)** Immunoblot showing CRR1 expression. Progeny of a tetratype tetrad (Supplemental Table 1) were grown in aerated minimal medium. Total cell proteins were processed using Immunoblot Analysis Method 1). The blot was probed with CRR1-B antibody (Supplementary Figure 2). An unidentified cross-reacting band at 72 kDa serves as a loading control. See Supplementary Figure 3 for the full-length immunoblot.

*CRR1* transcripts are equally abundant in both copper-replete (+Cu) and -Cu cells (Kropat et al. 2005). Abundance of the CRR1 protein was monitored using anti-CRR1 antibodies (see Methods; Supplementary Figures 2 and 3; Supplementary Results for Supplementary Figure 3). Immunoblot analysis indicates that the protein accumulates only in -Cu cells and is lost within minutes when Cu is added to the culture (Figure 9). The protein is present in wild-type but not *crr1* cells (Supplementary Figure 3C). Consistent with its proposed function, it is found in the nuclear fraction of cells grown in copper-deficient medium (Supplementary Figure 3D). We also conclude that levels of CRR1 protein are regulated at a post-transcriptional level.

**Figure 9.**
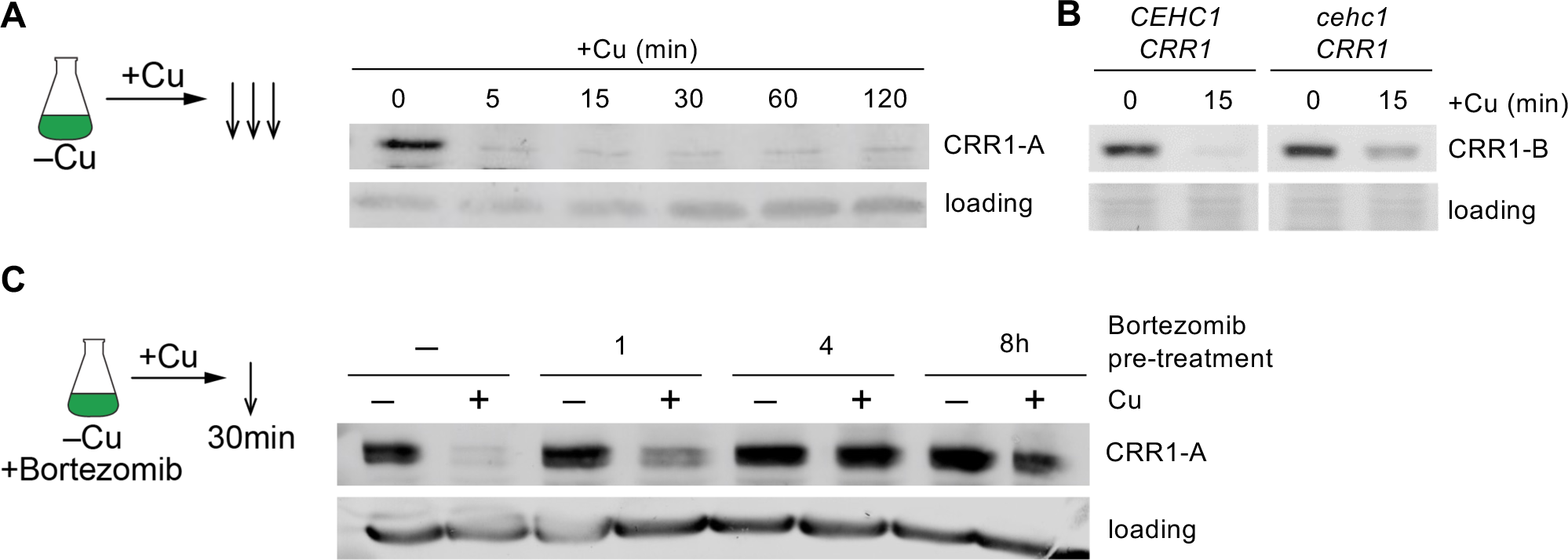
Pre-incubation with a proteasome inhibitor blocks Cu-dependent loss of CRR1. **A)** CuCl_2_ (2 μM) was added to copper-deficient cells and cells were sampled at the indicated times thereafter. Cells were re-suspended in buffer containing a protease inhibitor cocktail and processed as described for visualization of CRR1 abundance by immunoblot analysis (method 2) using CRR1-A antiserum. **B)** CuCl_2_ (2 μM) was added to Cu-deficient cells of the *CEHC1/CRR1* and the *cehc1/CRR1* genotypes of the tetratype tetrad (Supplemental Table 1) and cells were sampled at the indicated times thereafter. Cells were processed as described in methods for visualization of CRR1 abundance by immunoblot analysis using CRR1-B antiserum. **C)** Bortezomib was added to Cu-deficient cells to a final concentration of 20 μM from a stock solution of 10 mM in DMSO at time 0h. CuCl_2_ (2 μM) was added at 1, 4 and 8h after Bortezomib and the cells were sampled 30 min later for visualization of CRR1 by immunoblot analysis using the CRR1-A antiserum. Full blots are shown in Supplementary Figure 2.

An elevated level of CRR1 in copper-replete cells was observed only in cells with the *cehc1-1*; *CRR1* allele combination (Figure 8C), the same combination found in cells with elevated expression of the reporter gene and CRR1 target genes (Figure 8A, B). In contrast, cells with the *CEHC1* allele, including B6 and phenotypically rescued cells, have much lower levels of CRR1. Copper addition to -Cu wild-type cells resulted in CRR1 loss within minutes (Fig. 9A, B), whereas the same treatment of *cehc1-1; CRR1* cells resulted in partial CRR1 loss (Figure 9B). The intermediate level of CRR1 protein present in the *cehc1-1* mutant strain after copper addition may reflect a low level of CEHC1 protein remaining in *cehc1-1* cells caused by splicing flexibility (Figure 4C). The result is consistent with a function for CEHC1, a candidate component of an SCF ubiquitin ligase, in the pathway leading to CRR1 destabilization.

The role of proteasomes in Cu-dependent destabilization of CRR1 was tested by pre-treatment of -Cu cells with bortezomib, an inhibitor of the catalytic activity of the proteasome (Fig. 9C). Pre-treatment allowed CRR1 to accumulate after Cu addition, indicating that regulation by Cu of CRR1 function involves its rapid degradation by the proteasome.

### The *cehc1-1* mutation does not affect cell growth

To examine the possible role of CEHC1 in the regulation of cell growth, tetrad progeny containing all combinations of the wild-type and mutant alleles at the *CEHC1* and *CRR1* loci were grown in minimal medium in the light. In growth curves obtained for two different sets of tetrads, we observed no consistent differences among the tetrad progeny (Supplementary Figure 5). The results suggest that the *CEHC1* protein is not a regulator of cell growth, despite the large number of genes whose expression is up- or down-regulated in *cehc1-1* mutants.

## Discussion

Our genetic screen identified a negative regulator of the *HYDA1* expression pathway. Conditional motility dependent on hypoxia was provided by the *HYDA1-RSP3* reporter gene expressed in *pf14* cells. The selection of motile cells from among 10^9^ chemically-mutagenized immotile cells revealed mutants with constitutive activity of the *HYDA1* promoter even in aerobic conditions, allowing the cells to produce wild-type RSP3 protein. The *CEHC1* gene is not closely linked to the reporter gene construct, consistent with a defect in a trans-acting factor that negatively regulates the pathway, thereby leading to increased transcription of *HYDA1*.

The ability of the *RSP3* gene to induce motility when transformed into cells with cilia paralyzed by the *pf14* mutation provides a versatile tool to study Chlamydomonas gene promoters. A screen for motile cells simply requires that they swim to the top of a liquid culture, meaning that very large numbers of mutagenized cells can be screened for a desired phenotype by inoculating sufficient numbers of culture tubes. The *RSP3* reporter was used previously as a promoter trap (Haring and Beck, 1997) and to report the induction of both strong and weak promoters including those from the *PSAD1*, *CYC6*, and *CAH1* genes (Ferrante et al. 2011). In a fusion construct with the *NIT1* promoter, it was used to find mutations in regulatory genes controlling ammonia repression of *NIT1* gene expression (Zhang and Lefebvre, 1997). Because most signaling pathways utilize both positive and negative regulatory genes, results from this and previous studies demonstrate the potential of using the *RSP3* reporter gene to study other pathways.

The possible role of the CEHC1 protein in ubiquitylation is suggested by its homology with mammalian and plant proteins (Fig 5). A search of the Conserved Domain database at NCBI (Lu et al. 2020) identified three conserved domains: F-box-like (PF12937), SMI1/KNR4 (PF09346), and ApaG (PF04379). An F-box contains ∼50 amino acids forming a triple helix surface that functions in protein-protein interactions (Bai et al. 1996). Based on additional interaction domains found in CEHC1 (described below), it is included along with its plant and metazoan homologs in the FBXO3 (Fbxo3) class of F-box-like proteins (Jin et al. 2004).

The Smi1/Knr4 domain (PF09346) takes its name from a *Saccharomyces cerevisiae* protein termed a “hub” protein due to its physical and/or genetic interaction with ∼100 other proteins (Martin-Yken et al. 2016). Zhang et al. (2011) discovered that the Smi1/Knr4 domain is part of the SUKH domain superfamily consisting of five major families of proteins present in eukaryotes, bacteria and DNA viruses. Eukaryotic protein family members include Fbxo3 proteins and a subunit of tubulin polyglutamylase. They proposed that the SUKH domain serves as a scaffold for binding proteins for post-translational modification by glutamylation or ubiquitylation.

The ApaG or DUF525 (PF04379) domain was first described in the ApaG proteins from gram-negative bacteria (Farr et al. 1989). The domain is composed of seven antiparallel beta strands forming two beta sheets arranged in a sandwich configuration similar to the Immunoglobulin/Fibronectin III-type fold (Cicero et al. 2007). The ApaG domain is also present in a few eukaryotic proteins including FBXO3 and mammalian polymerase delta interacting protein (PDIP38), a multifunctional protein that binds DNA polymerase δ (Xie et al. 2005). The X-ray structure of the ∼125 amino acid ApaG domain from the human FBXO3 protein was superimposable on that of the bacterial proteins (Krzysiak et al. 2016).

The identities and arrangement of predicted structural domains in the CEHC1 protein are similar to those of human FBXO3 (NP-036307.2) (Ilyin et al. 2000; Jin et al. 2004), a protein that functions in the ubiquitin-proteasome system. When bound to a substrate, F-box proteins form a complex with SKP1, an adaptor for the CUL1-RBX1 class of Cullin-RING E3 ubiquitin ligases (Harper and Shulman, 2021). A Cullin protein scaffold together with a RING ("really interesting new gene") domain protein regulates transfer of the ubiquitin from a ubiquitin-carrying enzyme to the target protein (Cardozo and Pagano, 2004; Deshaies and Joazeiro, 2009; Buetow and Huang, 2016). A well-known example is the SCF (SKP1-CULLIN1/CDC53-F-box protein) complex involved in cell cycle control. It contains four key subunits: Cullin1 (CUL1 or CDC53 in budding yeast), the RING domain protein RBX1 (ROC1, HRT1), the bridging protein SKP1, and the F-box containing substrate recognition protein SKP2 recognized by SKP1. Through a variety of C-terminal domains, different F-box proteins provide specificity for recognition of phosphorylated target proteins.

Activities of mammalian FBXO3 provide insights into the possible role of the CEHC1 protein. The human protein FBXO3 co-immunoprecipitated with CUL1, SKP1, and RBX1, indicating that it is a component of a Cullin-based E3 ligase and likely functions in ubiquitylation (Shima et al. 2008). Murine and human FBXO3 play a role in transcriptional activation of proinflammatory cytokines in the innate immunity system when signals from microbial infection stimulate cell surface receptors including the tumor necrosis factor receptor (Mallampalli et al. 2013). Associated factors (TRAFS) transduce signals leading to increased transcription of cytokines that, when uncontrolled, constitute a “cytokine storm”. In resting cells, TRAFS are maintained at low levels as a result of their binding to FBXL2, a component of an SCF-type E3 ligase, leading to their polyubiquitylation and proteasomal degradation (Chen et al. 2013). Levels of FBXL2 itself are regulated by phosphorylation, followed by polyubiquitylation and proteasomal degradation. Among many F box proteins tested, only FBXO3 overexpression resulted in instability of FBXL2 (Chen et al. 2013). Further, co-immunoprecipitation experiments showed the interaction of FBXL2 and FBXO3. The ApaG/DUF25 domain of FBXO3 is essential for this interaction and the degradation of FBXL2 (Mallampalli et al. 2013). In addition to its role in promoting cytokine production, FBXO3 has a role in several other regulatory pathways (reviewed by Zhang et al. 2022). For example, (Lai et al. 2016) found that it functions in polyubiquitylation of FBX2 in presynaptic neuronal cells. Although it is a negative regulator of FBXL2 and FBX2, FBXO3 also acts as a positive regulator in certain pathways. In the thymus, FBXO3 binds phosphorylated AIRE, the autoimmune regulator transcription factor, leading to monoubiquitylation of AIRE. The modification promotes interaction with the transcription elongation factor b (P-TEFb), leading to increased transcription of AIRE-responsive genes needed for maintenance of self-tolerance (Shao et al. 2016). The similarity of *CEHC1* to metazoan *FBXO3* suggests a role for *CEHC1* in regulating transcription factors.

The CEHC1 protein shows greatest sequence conservation with proteins encoded in numerous genomes in the Viridiplantae lineage such as the *Arabidopsis thaliana* SKIP16 protein (SKP1/ASK-interacting protein 16, At1g06110) (Risseeuw et al. 2003; S1 Fig). SKIP16 interacts with SKP1-like proteins ASK2 (At5g42190) and ASK 4 (At120140) (Risseeuw et al. 2003) (Arabidopsis Interactive Mapping Consortium, 2011; BioGRID: https://thebiogrid.org) in yeast two-hybrid experiments. These results indicate that SKIP16 is likely a component of a RING domain E3 ligase and involved in a protein ubiquitylation pathway. Although SKIP16 transcripts are widely expressed in *A. thaliana* tissues (Klepikova et al. 2016), no function has yet been described for this gene.

If the CEHC1 protein functions similarly to FBXO3 homologs in other systems, it will bind a substrate protein (one of which we propose is CRR1) that becomes polyubiquitinated and then degraded by proteasomes, causing its cellular levels to fall. When CEHC1 protein function is lost through mutation, CRR1 would not be subject to degradation and its steady state levels would increase, resulting in increased expression of the CRR1 regulon, including increased transcription of HYDA1 in response to dark hypoxia. In aerobic conditions and in copper-supplemented cells, the CEHC1 protein likely functions to keep the levels of CRR1 low and thus acts as a negative regulator of the pathway. Mutation of *CEHC1* leads to constitutive activity of CRR1, increasing transcription of the p*HYDA1-RSP3* reporter gene, the endogenous *HYDA1* gene, and many others.

Our results indicate that *CEHC1* functions upstream of *CRR1*. The *crr1* mutant fails to grow in anoxia, even in copper-replete medium. Upregulation of transcript levels for certain indicator genes controlled by *CRR1*, including those encoding cytochrome *c*_6_ (*CYC6*), coproporphyrinogen oxidase (*CPX1*) and a di-iron enzyme (*CHL27A*, formerly *CRD1*), were found as common responses in both the copper-sensing and hypoxia-sensing pathways (Quinn et al. 2000, 2002). CRR1 binds the GTAC-containing copper response element in promoters of target genes via a *squamosa*-promoter binding protein (SBP) domain (Kropat et al. 2005; Sommer et al. 2010). A metallothionein-like Cys-rich domain at the C-terminus is required for the *CRR1* response to hypoxia (Sommer et al. 2010; Hemschemeier et al. 2013), suggesting a mechanism for CRR1 activity in the integration of copper response and anoxia response pathways. In a previous genetic screen for positive regulators of the *HYDA1* promoter in response to hypoxia, Pape et al. (2012) employed an arylsulfatase reporter gene with the *HYDA1* promoter and obtained a mutant in *CRR1*. Target genes for *CRR1* were identified by comparison of transcriptomes from wild-type cells with those of *crr1-2* mutant cells in response to copper deprivation or hypoxia and by comparison of transcriptomes from cells changing from copper replete to copper-depleted medium (Castruita et al. 2011; Hemschemeier et al. 2013; Kropat et al. 2015; Blaby-Haas et al. 2016). The *HYDA1* gene was found to be upregulated in response to hypoxia and in *crr1-2* mutant cells, although to a lesser extent than in wild-type cells (Pape et al. 2012; Hemschemeier et al. 2013; Blaby-Haas et al. 2016), suggesting that another mechanism(s) affects *HYDA1* transcript accumulation in addition to increased transcription driven by CRR1.

Given previous results showing a role for CRR1 in the signaling pathway for control of *HYDA1* expression, it is not surprising that our study uncovered genetic interaction between *CEHC1* and *CRR1*. Activity of *CRR1* is required for expression of the p*HYDA1-RSP3* reporter gene but this occurs only in the *cehc1-1* mutant (Fig 8A). Similarly, endogenous genes known to be targets of CRR1 are upregulated in the *cehc1-1*; *CRR1* background. The RNA-seq data showed that known target genes of *CRR1* are upregulated (or down-regulated) in the *cehc1-1* mutant and that this expression change is rescued in mutant cells transformed with the wild-type *CEHC1* gene. These data are consistent with a model in which *CEHC1* and *CRR1* act in the same signaling pathway and that the *CEHC1* gene acts upstream of *CRR1*. For purposes of discussion, we will assume that *CEHC1* acts on *CRR1* itself, although it could act on some other proteins required for *CRR1* function. The levels of *CRR1* transcripts do not change in the *cehc1-1* mutant, a result consistent with the observation that *CEHC1* regulates half-life of the CRR1 protein (Figure 9) rather than transcription or half-life of the mRNA. Loss of CEHC1 activity in the *cehc1-1* mutant would be expected to stabilize CRR1, leading to higher levels of *HYDA1* transcripts, as we observed for both the *HYDA1* endogenous gene and for the reporter gene driven by the *HYDA1* promoter. Using an inhibitor of proteasomes, bortezomib, we showed that CRR1 is subject to proteasomal degradation (Fig 9).

Among the questions raised by this study is whether *CEHC1* acts on other pathways in addition to *CRR1*. The RNA-seq results showed that in *cehc1-1* mutants, transcript levels were altered for genes not known to be regulated by *CRR1*. Future experiments could address this question by comparing RNA-seq results among cells with different allele combinations of the *CEHC1* and *CRR1* gene and exposed to different copper and hypoxia conditions. Investigating the possible physical interaction between the F-box domain of CEHC1 and the CRR1 protein would provide confirmation of the genetic interaction obtained in our study. The human homolog of CEHC1 protein, FBXO3, acts to degrade another F-box protein, FBXL2, which contains leucine-rich repeats. Although the Chlamydomonas genome contains a similar F-box protein with leucine-rich repeats (Cre01.g047650), it is not known whether it interacts with CEHC1.

Green algae like Chlamydomonas offer an attractive prospect for renewable biohydrogen production. However, two impediments must be overcome to allow the development of this potential energy source. Both the expression of hydrogenase genes and the activity of hydrogenase enzymes are tightly regulated by oxygen levels. Using a novel ciliary reporter gene system to identify mutants with constitutive expression from the *HYDA1* promoter we identified *CEHC1* as a negative regulator of *HYDA1* expression. Elimination of this negative regulator in the pathway for control of gene expression results in constitutive expression of transcripts encoding hydrogenases and related genes, even in the presence of normal oxygen levels. The *cehc1-1* mutant strain provides a starting point for efforts to engineer an oxygen-tolerant hydrogenase that could be expressed in the *cehc1-1* mutant background. In addition, the *cehc1-1* mutant could be used as a starting strain to select for natural mutations in hydrogenase genes that would produce oxygen-tolerant enzymes.

## Methods

### General Growth Conditions

Cells were maintained on Tris-Acetate-Phosphate (TAP) with Hutner’s trace elements or Sager-Granick (M) medium (Harris,1989) agar for long-term storage https://www.chlamycollection.org/methods/media-recipes/. Strains with the *ac17* mutation were maintained on TAP agar or ½ R agar (M medium with a three-fold increase in potassium phosphate and supplemented with 11 mM sodium acetate). Strains with the *arg2* mutation were kept on TAP-Arg (0.02 %) medium. Liquid cultures were grown in TAP or M media bubbled with filtered air, at 24°C on a 14-hr light/10-hr dark cycle, illuminated with white light (4800 lux) from fluorescent tubes. For experiments shown in Figure 9 and Supplementary Figure 3, cells were grown in TAP medium with revised trace elements (Kropat et al., 2011) and continuous illumination.

### Genetic Analysis

Gametogenesis, mating, and tetrad analysis was performed at 24° C using standard protocols. In crosses of motile CC-4509 cells with immotile CC-613 cells, the progeny were scored as PD (2 motile:2 immotile; NPD (4 immotile); TT (1 motile:3 immotile). Linkage was calculated using the formula 1/2T + 3NPD/(PD + NPD +T) (Harris, 1989). A ratio of 43:1:33 indicated that *CEHC1* is linked within 25 cM of the site of genomic insertion of the reporter gene construct (Sun et al. 2013).

### Molecular mapping PCR

Genomic DNA from tetrad progeny of the B6-F x CC-2290 cross was extracted using Puregene Core Kit (Qiagen 1042601). PCR reactions were carried out as described (Kathir et al. 2003) using primers from the molecular mapping kit (Chlamydomonas Resource Center, University of Minnesota; Kathir et al. 2003; Rymarquis et al. 2005) or primers designed for this project by BLAST comparison of DNA sequences from the reference genome (Phytozome v5.6; Merchant et al. 2007) with those from the polymorphic Chlamydomonas strain CC-2290 (Supplementary Tables 2 and 3).

### Generation and Analysis of Diploid Strains

Stable diploids were selected using the *nit1* mutation in the CC-4509 strain and the *ac17* mutation in strain CC-4322. The mating culture was plated on M medium lacking acetate and ammonium (substituting KNO_3_ for NH_4_NO_3_) to select for stable diploid *AC17 NIT1* colonies. After five days growth under constant light, colonies (potential stable diploid cells) were picked into liquid selective medium for analysis of motility using a stereomicroscope. To determine whether both mating type loci were present in a potential diploid line, DNA was isolated and used as a template for the PCR with primers for mt^+^ and mt^-^. Loss of heterozygosity for chromosome 1 was tested by using the PCR with primers designed (Supplementary Tables 4 and 5) to amplify DNA fragments containing SNP differences between the genome of the reference strain and the genome sequence of strain CC-4509, generated in this project. The amplified fragments were sequenced to determine the presence of one or both alleles at the SNP site.

### Cloning and tagging the *CEHC1* gene

A WT copy of gene Cre01.g005900 was cloned by screening a lambda phage library of WT Chlamydomonas DNA (Schnell and Lefebvre, 1993) with a hybridization probe consisting of a 500 bp fragment from the 3’ end amplified by using PCR. Restriction mapping of the resulting clones showed that phage 17-1 contained the complete gene. A HindIII/SpeI fragment extending from Cre01: 1029777 – 1024634 in Phytozome v5.6 genomic DNA sequence was cloned into pBlueScript KS (+) to create plasmid pCEHC1-5.1H/S. The plasmid was modified by inserting three copies of a sequence encoding the HA epitope into the SacI site located immediately downstream of the ATG start codon. In addition, a 322 bp HindIII fragment was cloned into the HindIII site to extend the promoter region. The final clone is designated p*CEHC1*-5.5H/S(1)3xHA5’. A selectable marker gene was added by digesting the plasmid with ClaI and KpnI and ligating it with a 1.7 kb fragment containing the aph7” gene from *Streptomyces hygroscopicus* (Berthold et al. 2002). This gene was amplified from plasmid pHyg3 by using a forward primer at the 5’ end modified to include a ClaI site and a reverse primer at the 3’ end that included the KpnI site. This plasmid was designated p*CEHC1*-5.5H/S(1)3xHA5’+Hyg3. Plasmids were digested with KpnI for use in transformation experiments.

### Chlamydomonas Transformation

Glass bead co-transformation with plasmid pSI103 (Sizova et al. 2001) and selection on paromomycin was carried out as described (Sun et al. 2013). For phenotypic rescue of *cehc1-1* mutants, CC-4509 cells were co-transformed with a lambda phage or plasmid construct containing the Cre01.g005900 gene (p*CEHC1*-5.5H/S(1)3xHA5’) together with the pHyg3 plasmid (Berthold et al. 2002). In other experiments, a plasmid containing both the HA-tagged Cre01.g005900 gene and the *aph*7” gene (p*CEHC1-*5.5H/S(1)3xHA5’+Hyg3) was used for transformation. The transformation mixture was plated on TAP medium containing 18 μg/ml Hygromycin B (Roche 10 843 555 001; Mannheim, Germany) using 0.9% agar to screen for colony morphology. Phenotypic rescue of the *cehc1-1* mutation should result in downregulated expression of the p*HYDA1-RSP3* reporter gene and loss of motility. Because colonies of motile cells spread to a larger diameter on soft agar as compared to colonies of immotile cells (Bloodgood, 1981), we picked colonies with a tightly packed morphology into liquid culture and assayed cell motility using phase contrast microscopy.

### Immunofluorescence Labeling

Localization of HA-tagged CEHC1 and tubulin in autolysin-treated cells was carried out as described in Silflow et al., 2001.

### qPCR Quantification Using Roche Universal Probe Library

Assays were carried out at the Biomedical Genomic Center (BMGC), University of Minnesota. The assay design utilized the Roche Universal ProbeLibrary (UPL) technology with hydrolysis probes and was based on the JGI v4.0 *C. reinhardtii* genomic DNA sequence (http://genome.jgi.doe.gov/Chlre4/Chlre4.home.html). Each assay design generated a sequence for the forward primer, reverse primer, amplification product (amplicon) and provided the UPL probe number. In most cases, the primer sets spanned an intron (Supplementary Table 8). To generate cDNA for the assays, total RNA samples were treated with DNase using the TURBO DNA-free kit (Applied Biosystems). The RNA was used as a template for first-strand cDNA synthesis using SuperScript II reverse transcriptase (Invitrogen) following the protocol of the manufacturer and using an Applied Biosystems GeneAmp PCR System 9700.

To validate the primer probe set designs, a one-to-five dilution series of the cDNA sample was created in five wells of a 96-well plate with the starting concentration at 5 ng/μl. Four μl of a cocktail containing 2 μl of 10 mM forward primer, 2 μl of 10 mM reverse primer, 1 μl of 10 mM probe (Roche Universal ProbeLibrary), and 36 μl of 2.2X Master mix was dispensed into wells containing 4 μl aliquots of the dilution series or a 4 μl aliquot of distilled water for a negative template control. 3 μl of the dilution series and cocktail mix were dispensed into duplicate wells on a 384-well ABI optical plate and the plate was loaded on the ABI 7900HT Fast Real-Time PCR System (Applied Biosystems). The FAM-Non Fluorescent Quencher detector and Rox passive reference were selected. The qPCR protocol was: 2 min activation at 60° C, 5 min denaturation at 95° C, followed by 45 cycles of 10 sec at 95° C and 1 min at 60° C. Different iterations of the primer probe set designs were tested with the dilution series. The set design closest to 10% efficient was chosen for the expression run. Any design <90% efficient or >110% efficient was rejected.

For quantitative PCR reactions, 3 μl of cDNA (24 ng) was dispensed into duplicate wells on a 384-well ABI optical plate. A working cocktail contained 10 μM forward primer (IDT), 10 μM reverse primer, 10 μM probe (Roche Universal ProbeLibrary), and 2.2X Master mix. 3 μl of cocktail was dispensed into the wells containing the cDNA samples and the reactions were carried out as described above.

The results of qPCR are presented as Ct (threshold cycle) values. Data were analyzed using the ΔΔCt method (Livak and Schmittgen, 2001). All data were normalized to the transcript levels of RCK1 (Cre06.g278222; Schloss,1990) and calibrated to the transcript levels of a control strain.

### Reverse Transcription PCR

Total RNA (1 μg) was treated with DNase I (Amplification grade, Invitrogen) at room temperature for 45 min and the enzyme was inactivated as per instructions of the manufacturer. First-strand cDNA synthesis was carried out with SuperScript III reverse transcriptase (Invitrogen) following the manufacturer’s protocol in a total volume of 50 μl. The RT primer (5 pmol, Supplementary Table 7) was used to initiate reverse transcription. A 5 μl aliquot of the reverse transcription product was used as the template for PCR amplification in a total volume of 50 μl, using FailSafe enzyme mix and FailSafe PreMix K from Epicenter Biotechnologies (Madison, WI) along with gene-specific primers (Supplemental Table 7). Amplification was performed following standard cycles: 95° C 3 min, 35 cycles of (95° C 30 sec, 53° C 30 sec, 72° C 30 sec), 72° C 5 min.

### Next-generation Whole Genome Sequencing

Genomic DNA samples prepared using the CsCl centrifugation method (Schnell and Lefebvre, 1993) from strains CC-4507 and CC-4509 were used for whole genome next-generation sequencing. The sequencing libraries were prepared by the BMGC at the University of Minnesota, using Nextera Library Prep Kits (Illumina), following the manufacturer’s instructions. Illumina cBOT (Illumina) was used for cluster generation, following the manufacturer’s instruction. Both DNA samples were sequenced for paired-end, 100-bp read length using the Illumina GAIIX sequencer. Short reads were trimmed for a minimum quality score of 28 at the 3’ end. A total of ∼20.9 million pass-filter reads (274 bp average) were generated for strain B6 and ∼27.6 million (320 bp average) for strain B6-F. The data provided a roughly 24-fold coverage for the 120 Mb nuclear genome (Merchant et al. 2007) of strain B6 and 37-fold coverage for strain B6-F. Reads were aligned to the *C. reinhardtii* v5.6 genome at JGI using BWA (Burrows-Wheeler Aligner) (Li and Durbin, 2009) and Bowtie (Langmead et al. 2009) with default settings. SNPs were identified by comparing uniquely mapped reads generated from Bowtie alignment between the two DNA samples. SNP sites with a coverage of less than 6 reads for either strain were excluded from further analysis. For cases of multiple base calls, the required allele ratio at the position was >85%. After filtering, a total of 49 SNPs were identified between the genomes.

### Preparation of CRR1 antibodies

#### CRR1-A

A portion of CRR1 (439-800 including the SBP domain and the extended SBP domain, Epitope 3, Supplementary Figure 2) was expressed in *Escherichia coli* with an amino-terminal his_6_-tag and purified in 50 mM Tris-Cl pH 8, 100 mM NaCl, 10 mM 2-mercaptoethanol, 6 M urea at the UCLA Protein Expression facility. The purified protein was used to immunize rabbits at Covance (now Labcorp), Denver, PA. The purified antigen was also used by Covance to affinity purify antibodies from pooled serum generated from the immunization protocol.

#### CRR1-B

A mixture of peptides corresponding to residues 1-21 (Epitope 1) and 34-50 (Epitope 2), conjugated to keyhole limpet hemocyanin, were used to immunize rabbits at Labcorp, Denver, PA. Antibodies in the resulting serum were affinity purified using the same peptides. Validation of the antibodies is presented in Supplementary Figure 3 and discussed in the Supplementary Results.

### Immunoblot Analysis

#### Method 1

Protein extracts from intact cells were prepared by pelleting 1.8×10^7^ cells, removing the supernatant, adding 180 μl of 1.5x sample buffer (93.75 mM Tris-Cl, pH 6.8, 3% SDS, 15% glycerol, 7.5% 2-mercaptoethanol), and immediately boiling the mixture for 90s. For detection of CRR1, concentrated cell culture was resuspended in M medium containing 1X protease inhibitor cocktail before boiling (Sigma P8340). Before loading the gel, the sample tubes were centrifuged at 14,000g for 2 min to pellet cell debris and gel lanes were loaded with 20 ul of the solution (1.8 × 10^6^ cells per lane). Proteins were fractionated using SE250 Mighty Small II (Hoefer Scientific Instruments) on SDS-PAGE minigels (1.5 mM thick, 8% polyacrylamide) and transferred to Immobilon-P membranes (Millipore IPVH00010) using a Bio-Rad Mini Trans-Blot (#1703930) with 0.192 M Tris-Glycine buffer (pH 8.3) in 20% methanol at 102 V for 70 min. HA-tagged proteins were detected using the ECL Western Blotting Analysis System (Amersham #RPN2109) protocol. Blots were incubated for three hours in anti-HA High Affinity antibody from rat IgG (clone 3F10; Roche), used at 1:1200 dilution in TBS Tween 20 (0.1%, pH7.6) containing 2.5% (w/v) dried milk. The secondary antibody was rabbit anti-rat IgG conjugated to peroxidase (Sigma-Aldrich A5795), used at a 1:8000 dilution in TBS-T with 2.5% dried milk buffer for one hour. For detection of CRR1, the primary antibody was CRR1-B at 1:1000 dilution. Secondary antibody was GAR-POD (Jackson Immunoresearch Lab., Inc. 111-035-144) at 1:50,000. After washing, blots were incubated in detection solutions 1 and 2 (ECL kit) and exposed to Blue Ultra Autorad film (GeneMate # F-9029-8X10). As a loading control, we used a rabbit antibody against OEE1 (gift from the Merchant laboratory) at a 1:3000 dilution. The secondary antibody was goat anti-rabbit horseradish peroxidase conjugate (Chemicon International AP183P) at a 1:22,000 dilution. Prestained molecular weight standards were from Thermo Scientific (Spectra Multicolor High Range Protein Ladder #26625). Proteins in the gel were silver-stained (Wray et al. 1981).

#### Method 2

For analysis of total proteins, 15 ml of a culture (density between 4-8 × 10^6^ cells/ml) was centrifuged at 1650*g*. The resulting cell pellet was resuspended in 300 μl of a buffer composed of 10mM Na-phosphate (pH 7), and EDTA-free cOmplete Protease Inhibitor Cocktail (Roche). Samples were flash frozen in liquid nitrogen and stored at -80°c prior to analysis. Protein amounts were determined using a Pierce BCA Protein Assay Kit against BSA as a standard and diluted with 2x sample butter (125 mM Tris-HCl pH 6.8, 20% glycerol, 4% SDS, 10% β-mercaptoethanol, 0.0005% bromphenolblue). Proteins were separated on SDS-containing 10% polyacrylamide gels using 10 μg of protein for each lane. The separated proteins were transferred by semi-dry electro-blotting to nitrocellulose membranes (Amersham Protran 0.1 or 0.45 micron NC as indicated in the legend, Supplemental Figure 3). The membrane was blocked for 30 min with 3-5% dried non-fat milk in phosphate buffered saline (PBS) solution containing 0.1% (w/v) Tween 20 (PBST) and incubated in primary antiserum. The PBS solution was used as the diluent for both primary and secondary antibodies. The membranes were washed in PBS containing 0.1% (w/v) Tween 20. Antibodies directed against CRR1 (1:1000 dilution), histone H3 (Abcam ab1791; 1:1000 dilution), COXIIb (Agrisesra AS06 151 at 1:4000 dilution) and PsaF (gift from J.D. Rochaix; 1:2000 dilution). The secondary antibody was goat anti-rabbit conjugated to alkaline phosphatase (ThermoFisher, Cat # 31340, 1:5000) and processed according to the manufacturer’s instructions.

### RNA-seq Analysis

Strains were grown in triplicate cultures in TAP medium under a 14h:10h light/dark regimen at 24°C. Light at ∼146 μmol m^-2^s^-1^ was provided by two 4-ft fluorescent 40W cool white bulbs placed over the cultures and two 4-ft LED 42W 3700 luminous flux lights placed to the side of the cultures. The cells were grown first in 10 ml cultures in 50 ml culture tubes to a density of 5 x10^6^ cells/ml. The cultures were transferred to 300 ml contained in 500 ml flasks and aerated with air bubbled through Pasteur pipettes until the cell density reached ∼5 × 10^5^ cells/ml. Finally, cells were diluted to 2.5 × 10^5^ cells/ml in 200 ml of fresh medium in 2800 ml Fernbach flasks (Pyrex 4420). The cultures were aerated by rotary shaking at 95 rpm for two days at which time the cell concentration was ∼5 × 10^6^ cells/ml. At 6 h into the light cycle, an aliquot of 3 × 10^7^ cells was removed from each flask and placed in a 15 ml conical tube. Processing each cell sample took approximately 1.5 min during which time the cells were pelleted by centrifugation for 45 sec, the supernatant was decanted, and the cells were suspended in 3 ml lysis buffer. Total RNA was prepared using the LiCl precipitation method (Wilkerson et al. 1994). The RNA samples were subjected to quality control tests using a Ribo Green Assay (Invitrogen) for quantification and an Agilent Nano chip (Agilent Technologies) to verify the RNA integrity. All samples surpassed an RNA Integrity Number of 8. A cDNA library was prepared from each RNA sample using the Illumina TruSeq RNA library preparation kit (Illumina) following the manufacturer’s instructions. The libraries were gel size selected to have average inserts of ∼200 bp.

The pooled libraries were sequenced in one lane on a HiSeq 2500 instrument using v4 chemistry and a 125 bp paired-end run. The lane generated ∼220 million reads with all expected barcodes well represented. Average quality scores were above Q30 for all pass-filter reads. Illumina conversion software bcl2fasstq version 2.17.1.14 was used to demultiplex and generate FASTQ files. To trim the sequences, Trimmomatic version 0.33 was used with parameters LEADING:3, TRAILING:3, SLIDING WINDOW:4:16, MINLEN:63 (Bolger et al. 2014). Average size of reads was 126 bp after trimming.

RNA-seq reads from strains B6 (*CEHC1*), B6-F (*cehc1-1*) and B6-F_resc (*CEHC1* rescued) were mapped to the *C. reinhardtii* reference assembly (v6 assembly, v6.1 annotations, available from https://phytozome.jgi.doe.gov) with RNA STAR (v2.7.10b; Dobin et al. 2013) with the following options: --alignIntronMax 3000 –outMultimapperOrder Random –outSAMmultNmax 1. Normalized counts for all nucleus-encoded transcripts were calculated in terms of fragments per kb of transcript per million mapped reads (FPKMs) with cuffdiff (v2.0.2; Trapnell et al. 2013) with the following options: --multi-read-correct –max-bundle-frags 1000000000 –library-type fr-firststrand. Reproducibility within each set of triplicate samples was indicated by the correlation coefficient, with average R values greater that .999 for each set (Supplementary Table 9). Differential expression analysis was performed in R with the DESeq2 package (v1.38.3). Genes were identified as differentially expressed (DEGs) if they met the following four criteria: 1) ≥5 FPKMs in at least one strain, 2) ≥ five-fold change in FPKMs between strains B6 and B6-F, 3) a Benjamini-Hochberg adjusted *p*-value < 0.05 for the fold change between strains B6 and B6-F, and 4) at least 50% rescue of expression in the B6-F_resc strain (calculated as the difference in FPKMs between B6-F and B6-F_resc divided by the difference between B6-F and B6).

In order to facilitate comparison of the *CEHC1*-regulated transcriptome with that of *CRR1*-regulated transcriptome, RNA-seq data from two previous studies was reanalyzed in parallel with this study using the same reference assembly and annotations. The first study examined the relationship between *CRR1* and Cu-responsive genes (*crr1-2* +/–Cu) (Castruita et al. 2011), and the second study examined the relationship between *CRR1* and dark anoxia-responsive genes (*crr1-2* +/–O_2_) (Hemschemeier et al. 2013). Raw sequencing data from both studies were downloaded from NCBI SRA from the following accessions: (PRJNA134525 for *crr1-2* +/–Cu and PRJNA178958 for *crr1-2* +/–O_2_), and mapped to the *C. reinhardtii* v6 assembly as described above. DEGs were identified using the same criteria as was reported in the previous two studies, respectively. For the Cu study, the inclusion criteria were: 1) ≥10 FPKMs in at least one strain, 2) ≥ two-fold change in FPKMs from the *CRR1* strain between –Cu and +Cu, and 3) ≥ two-fold change in FPKMs between the *crr1-2* mutant strain the *CRR1* complemented strain in –Cu. For the O_2_ study, inclusion criteria were: 1) ≥5 FPKMs in at least one strain, 2) ≥ two-fold change in FPKMs in the *CRR1* complemented strain between 6 h dark-anoxic and aerobic conditions, 3) <25% as much fold change in the *crr1-2* mutant strain between dark-anoxic and aerobic conditions relative to the *CRR1* complemented strain, and 4) a Benjamini-Hochberg adjusted *p*-value < 0.05 for the fold change in FPKMs in the *CRR1* complemented strain between 6 h dark-anoxic and aerobic conditions. A Venn diagram summarizing these results was produced with the VennDiagram package (v1.7.3) in R. All RNA-Seq data, including raw sequencing reads and normalized expression estimates, are available at the US National Center for Biotechnology Information (NCBI) Gene Expression Omnibus (GEO) repository under accession number GSE252595

## Supplementary Tables, Figures, Methods and Datasets

Supplementary Table 1. Chlamydomonas strains used in this study

Supplementary Table 2. SNP mapping of the *cehc1-1* mutation in strain B6-F

Supplementary Table 3. Primers for chromosome 1 mapping

Supplementary Table 4. Dominance/recessiveness testing of the *cehc1-1* mutation in strain B6-F

Supplementary Table 5. Primers for SNP markers developed in this study

Supplementary Table 6. Transformation rescue of the *cehc1-1* mutation in strain B6-F

Supplementary Table 7. Primers for Reverse Transcription (RT)-PCR

Supplementary Table 8. Primer/probe sets used in qPCR assays

Supplementary Table 9. Correlation Coefficients for replicate samples in RNAseq analysis

Supplementary Table 10. RNA-seq results for selected genes

Supplementary Method 1. Generation of a *crr1* mutant using CRISPR/Cpf1 gene editing

Supplementary Method 2. Isolation of Chlamydomonas nuclei

Supplementary Figure 1. Conservation of Fbxo3-like proteins in plants

Supplementary Figure 2. CRR1 amino acid sequence, domain structure, and epitopes.

Supplementary Figure 3 and Supplementary Results. Validation of CRR1 antibodies and full size CRR1 immunoblots and controls from Figure 9

Supplementary Figure 4. Full size Immunoblot of CRR1 from Figure 8

Supplementary Figure 5. Effect of *CEHC1*, *CRR1* mutant and wild-type alleles on cell growth

Supplementary Dataset 1. Combined FPKMs for All Genes

Supplementary Dataset 2. All 2FC DEGs in cehc1-1

Supplementary Dataset 3. Overlap of cehc1-1 and O2 Regulated DEGs

Supplementary Dataset 4. Overlap of cehc1-1 and crr1-2 DEGs

## Acknowledgments

Strains and plasmids were obtained from the Chlamydomonas Resource Center (CRC), supported by NSF grant 2247108. Strains and plasmids developed in this project were deposited in the CRC. We acknowledge Susan Perera for isolation of the SP0106B6 mutant strain (CC-4507), Paul Ranum for help with immunofluorescence localization. We thank the University of Minnesota Genomics Center for qPCR, RNA-seq, and DNA sequencing. Zheng Jin Tu and Juan Abrahante (UMGC) provided informatics support. We thank Mark Arbing at the UCLA Protein Expression Facility for preparing the recombinant his-tagged domain of CRR1. Financial support was provided by grants from the Office of the Vice-President For Research and the Institute for Energy and the Environment of the University of Minnesota (to CS). Supported by a grant from NIH, GM42143 (to SM).

## Author Contributions

Sun: designed the research, performed research, contributed new tools, analyzed data, wrote the paper

LaVoie: performed research, contributed new tools, analyzed data

Lefebvre: designed the research, wrote the paper

Gallaher contributed new analytic/computational tools, analyzed data, wrote the paper

Glaesener: performed research, contributed new tools

Strenkert: performed research, contributed new tools

Mehta: ( performed research

Merchant: designed the research, contributed new tools, analyzed data, wrote the paper

Silflow: designed the research, performed research, contributed new tools, analyzed data, wrote the manuscript

## Supplementary Figures, Results, and Methods for Sun et al

**Supplementary Figure 1.**
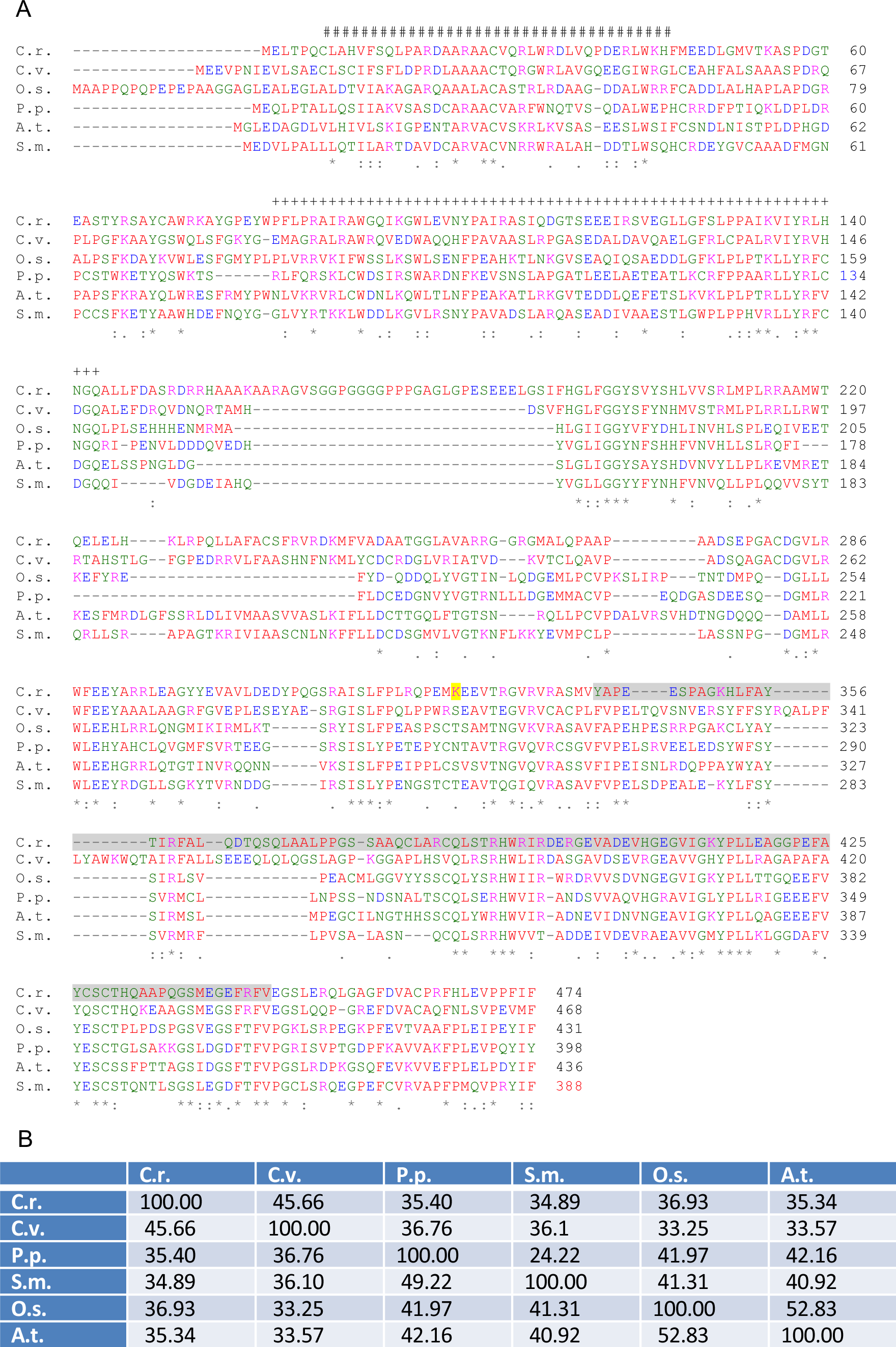
Conservation of FBXO3 proteins in plants. **A)** Amino acid sequences were aligned using Clustal Omega Multiple Sequence Alignment tools (Sievers and Higgins, 2021; EMBL-EBI). Accession numbers for the proteins are: C.r. (*Chlamydomonas reinhardtii* PNW87886.1, Cre 01.g005900, CEHC1); C.v. (*Chlorella variabilis* XP_005845474.1); A.t. (*Arabidopsis thaliana* NP_563759.1 SKP1/ASK-interacting protein 16); O.s. (*Oryza sativa* Japonica group NP_001067658.2); S.m. (*Selaginella moellendorffii* XP_002979125.1); P.p. (*Physcomitrella patens* XP_024376803.1). Domains for the *C. reinhardtii* protein were analyzed using the Conserved Domain Database at NCBI and SPARCLE software (Lu et al., 2020). The F-Box domain is indicated with # signs, the SMI1/KNR4 (SUKH-1) domain with + marks, and the DUF25 (ApaG) domain in gray text highlight. B. Percent Identity Matrix created by Clustal12.1 software (EMBL-EBI)

**Supplementary Figure 2.**
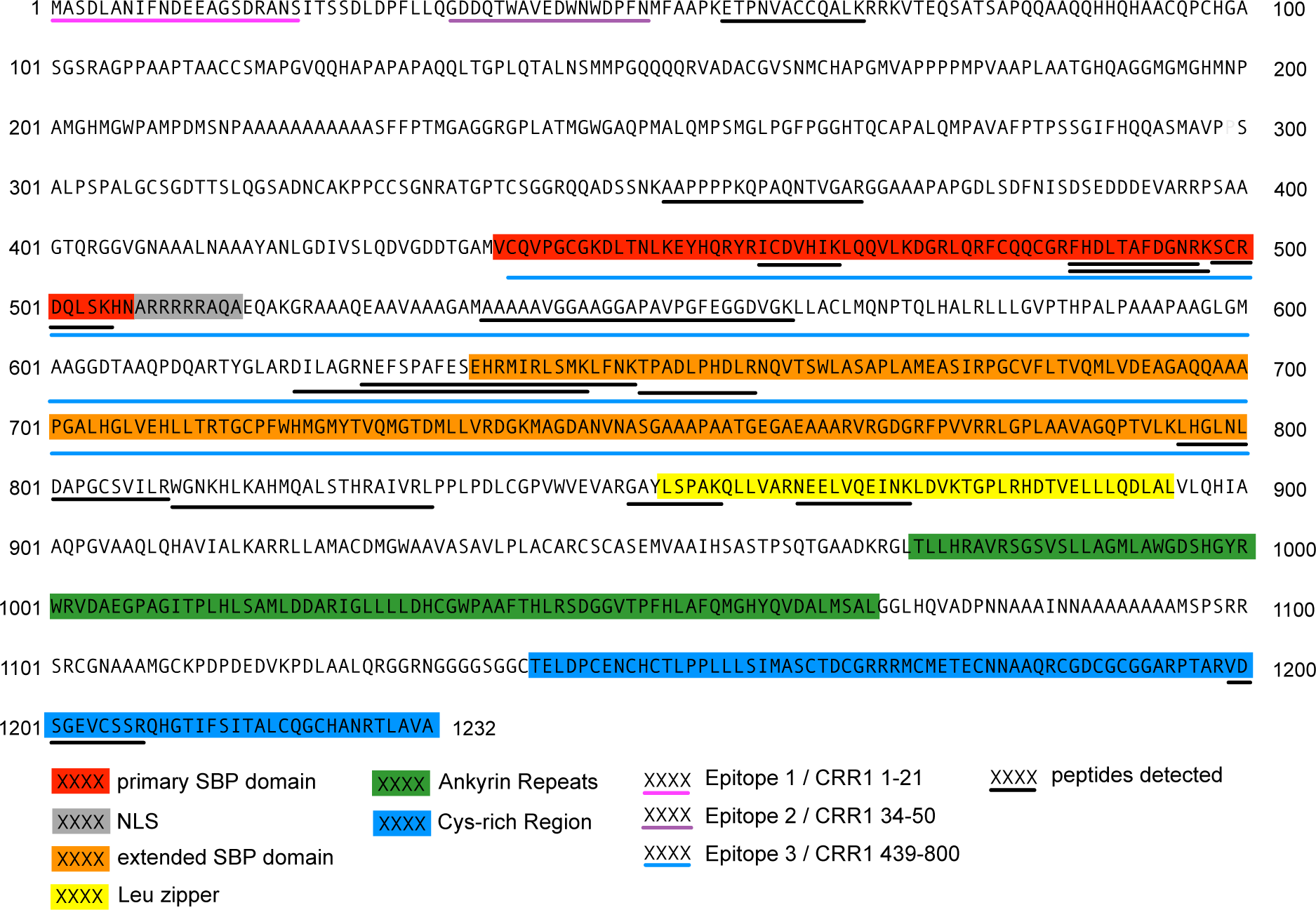
CRR1 amino acid sequence, domain structure, and epitopes. A peptide including amino acids 439-800 (Epitope 3) was used to raise antibody CRR1-A. A mixture of peptides corresponding to residues 1-21 (Epitope 1) and 34-50 (Epitope 2) was used to raise antibody CRR1-B. See Methods.

**Supplementary Figure 3.**
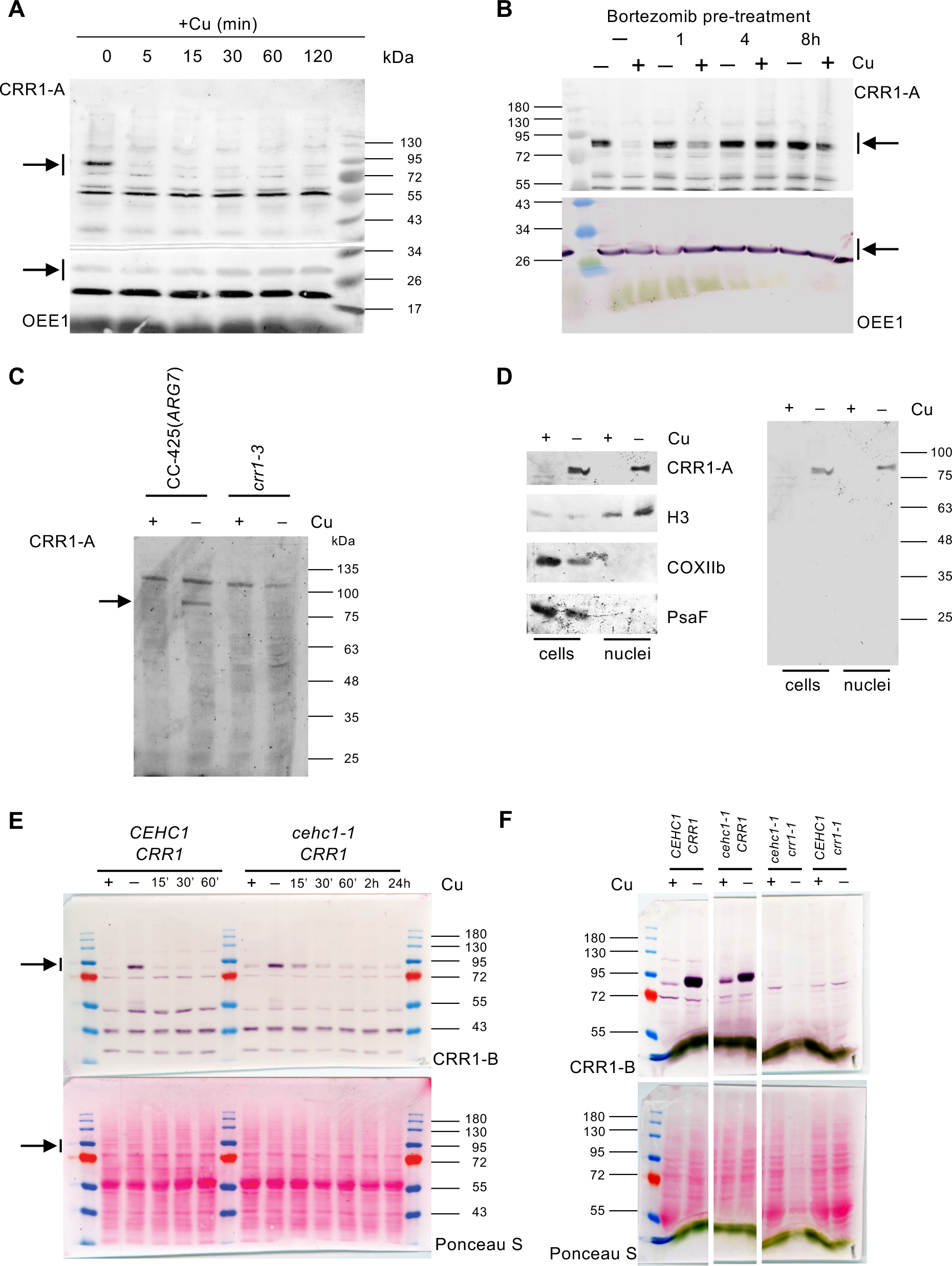
Validation of CRR1 antibodies and full size CRR1 immunoblots and controls. For all immunoblots in this figure, cells were re-suspended in buffer containing a protease inhibitor cocktail and processed as described in Protein Gels and Immunoblotting Method 2. **A)** Full size immunoblot, corresponding to Figure 9A. Copper was added to copper-deficient cells of strain CC-4532 and cells were sampled at the indicated times thereafter. The membrane (0.1 micron) was cut prior to immunodetection. Affinity purified CRR1-A antibody was used in the top section of the membrane and OEE1 antibody was used in the bottom section serving as loading control. The arrows to the left of the membranes indicate the section of the immunoblot shown in Figure 9A. **B)** Full size immunoblot, corresponding to Figure 9C. Bortezomib was added to Cu-deficient cells of strain CC-4532 at time 0h. Copper was added at 1, 4 and 8h after Bortezomib and the cells were sampled 30 min later for visualization of CRR1 by immunoblot analysis using the CRR1-A antiserum. The membrane (0.1 micron) was cut prior to immunodetection. Affinity purified CRR1-A antibody was used in the top section of the membrane and OEE1 was used in the bottom section serving as loading control. The arrows to the right of the membranes indicate the section of the immunoblot shown in Figure 9C. **C)** Immunodetection of CRR1 in *CRR1* and *crr1-3* cells using antibody CRR1-A. Arrow indicates CRR1 protein. **D)** Immunodetection of CRR1 in Cu-replete and Cu-depleted cells of strain CC-425. Whole cell extracts as well as a cell fraction enriched for nuclei are shown. Anti-histone H3 (H3) serves as a marker for nuclei, COXIIb for mitochondrial proteins and PsaF for chloroplast proteins. CRR1 is detected in both the whole cell extract as well as in the fraction highly enriched for proteins localized in the nucleus. **E)** Full size immunoblot, corresponding to Fig 9B. Cu was added to Cu-deficient cells of the *CEHC1/CRR1* and the *cehc1-1/CRR1* genotypes of the tetratype tetrad (Supplemental Table 1) and cells were sampled at the indicated times thereafter. Immunoblot analysis (0.45 micron nitrocellulose membrane) utilized CRR1-B antiserum. The arrows to the left of the membranes indicate the section of the immunoblot shown in Fig 9B. **F)** Full size immunoblots of *CRR1* and *crr1* cells using the CRR1-B antiserum. Cu replete and Cu deplete cells of the tetratype tetrad (Supplemental Table 1) were sampled at the indicated times thereafter. Immunoblot analysis used CRR1-B antiserum.

**Supplementary Results for Supplementary Figure 3**

Affinity purified anti-CRR1-A recognized a polypeptide that migrated in the range of 85 to 90 kDa (calculated from three different membranes) in whole cell extracts from Cu-deficient cells but not Cu-replete cells (Supplemental Figure 3C). The signal was absent in a *crr1* null mutant (*crr1-3*; generated by CRISPR, see Supplemental Methods). In previous work, the protein was suggested to be localized to the nucleus based on the occurrence of a nuclear localization signal and the localization of a GFP-CRR1 construct in mustard seedlings (Sommer et al., 2010). When we tested preparations of nuclei from Chlamydomonas, we noted that the immunoreactive band co-fractionated with histones, which further validated the authenticity of the signal. In many experiments, other signals were observed even with the affinity purified antibody (e.g. full-length panel in Supplemental Figure 3C), but these signals were found also in the null *crr1* strain, indicating that they are not specific to CRR1. An independent antiserum generated against two N-terminal epitopes, anti-CRR1-B, also recognized a similar sized protein (Supplemental Figures 3E, F), which was also present only in Cu-deficient cells (Supplemental Figure 3E) and in the wild-type CRR1 strain (Supplemental Figure 3F). Other bands were judged to be non-specific and these non-specific bands were different from those observed in blots probed with anti-CRR1-A. Based on the sequence of the ORF in the *CRR1* mRNA, we expected a protein of about 127 kDa. It is possible that the protein is either proteolytically processed after translation or it simply migrates faster than expected in denaturing gels. When we surveyed pooled Chlamydomonas proteomic data, we identified peptides along the length of the full protein (Supplemental Figure 2); therefore, we favor the latter hypothesis.

**Supplementary Method 1. Generation of a *crr1* mutant using CRISPR/Cpf1 gene editing**

CC425, a cell wall reduced arginine auxotrophic strain, was used for transformation with a RNP complex consisting of a gRNA targeting exon2 of CRR1 (Cre09.g390023) and LbCpf1 exactly as described in Pfam et al. (2022). In brief, 2 × 10^7^ cells were suspended and centrifuged (5 min, 1,500*g*) in Max Efficiency Transformation Reagent (1 mL) twice, followed by suspension in 230 μL of the same reagent supplemented with sucrose (40 mM). Cells were incubated at 40°C for 20 min. Purified LbCpf1 (80 μM) was preincubated with gRNA (2 nmol, TTTGGGGCTTGACATGC) at 25 °C for 20 min to form RNP complexes. For transformation, 230 μL cell culture (5 × 10^5^ cells) was supplemented with sucrose (40 mM) and mixed with preincubated RNPs and HinDIII digested pMS666 containing the *ARG7* gene conferring the ability to grow without arginine. In order to achieve template DNA-mediated editing, ssODN (∼4 nmol, sequence containing three in-frame stop codons after the PAM target site (CTGCATGGACGTTGGTAATCCGTAAGCGTTTTGGGGCTTGACATGCAGGCGACGATTAAACATAGGCCGTCGAGGACTGGAACTGGGACC) was added. Final volume was 280 μL. Cells were electroporated in a 4-mm gap cuvette (Biorad) at 600 V, 50 μF, 200 Ω by using Gene Pulser Xcell (Bio-Rad). Immediately after electroporation, cells were transferred and recovered overnight in darkness and without shaking in 4 mL TAP with 40 mM sucrose and polyethylene glycol 8000 0.4% (w/v) and then plated after a 5 min centrifugation at RT and 1650*g* using the starch embedding method. We used 60% starch as described in the following: Corn starch was washed sequentially twice with distilled water and once with ethanol. The washed starch was stored in 75% ethanol to prevent bacterial contamination. Before each experiment, the ethanol was replaced with TAP-sucrose medium by repeated centrifugations and resuspensions (5 times). The starch was finally resuspended to 60% (w/v) in TAP-sucrose medium and polyethylene glycol 8000 0.4% (w/v); 0.5 ml of the starch suspension was spread with one electroporation reaction after overnight recovery over the top of solid TAP medium in a petri dish plate. After 14 days, colonies were transferred to new plates and screened based on their inability to survive on TAP plates without Cu supplementation. gDNA from two candidate *crr1* lines was extracted using the CTAB method and PCR was performed using oligos Crr1seqfor: CAACTGCATGGACGTTGGTA and Crr1seqrev: GCTGGTGGTGCTGCTGTG. We used the BioRad iTaq SYBR green MasterMix and the following program for all qPCR reactions: 95°C for 5 min followed by 40 cycles of 95°C for 15 s, and 65°C for 60 s. Sanger sequencing of the PCR products using oligo Crr1seqrev revealed that ssODN mediated gene editing was not successful but showed a frame-shift in *crr1-3* most likely due to errors during DNA damage repair. The *crr1-3* mutant was used in experiments in Supplementary Figure 3.

**Supplementary Method 2. Isolation of Chlamydomonas Nuclei**

Nuclei were isolated from *Chlamydomonas* CC-5390 by a modified protocol described in Winck et al. (2011). In brief, a cell pellet (∼10^8^ cells) was resuspended in 1 mL of nuclei isolation buffer with protease inhibitors (NIBA) (1 mM phenylmethylsulfonyl fluoride (PMSF), 5 mM dithiothreitol (DTT), 10 mM sodium butyrate, and 1× protease inhibitor cocktail complete, without EDTA (Roche)). The resuspended cell pellet was added dropwise to a liquid-nitrogen-cooled mortar and ground to a fine, green powder. The sample was transferred to a 50 mL conical flask which was then filled to 20 mL with NIBA and the sample was centrifuged at 1260 *g* for 30 min at 4°C. The supernatant was discarded and the cell pellet was resuspended in 20 mL of NIBA + 1% Triton X-100 and mechanically lysed by pipetting ∼20–30 times with a Pasteur pipette. The sample was placed on ice for 10 min and centrifuged at 1000*g* for 30 min at 4 °C to collect nuclei. The supernatant was removed and mechanical lysis with NIBA + 1% Triton X-100 was repeated once again to ensure complete lysis of the plasma membrane. The nuclei pellet was transferred to an Eppendorf microcentrifuge tube, washed twice with 1 mL of NIBA to remove the detergent.This method was used for experiments in Supplementary Figure 3.

**Supplementary Figure 4.**
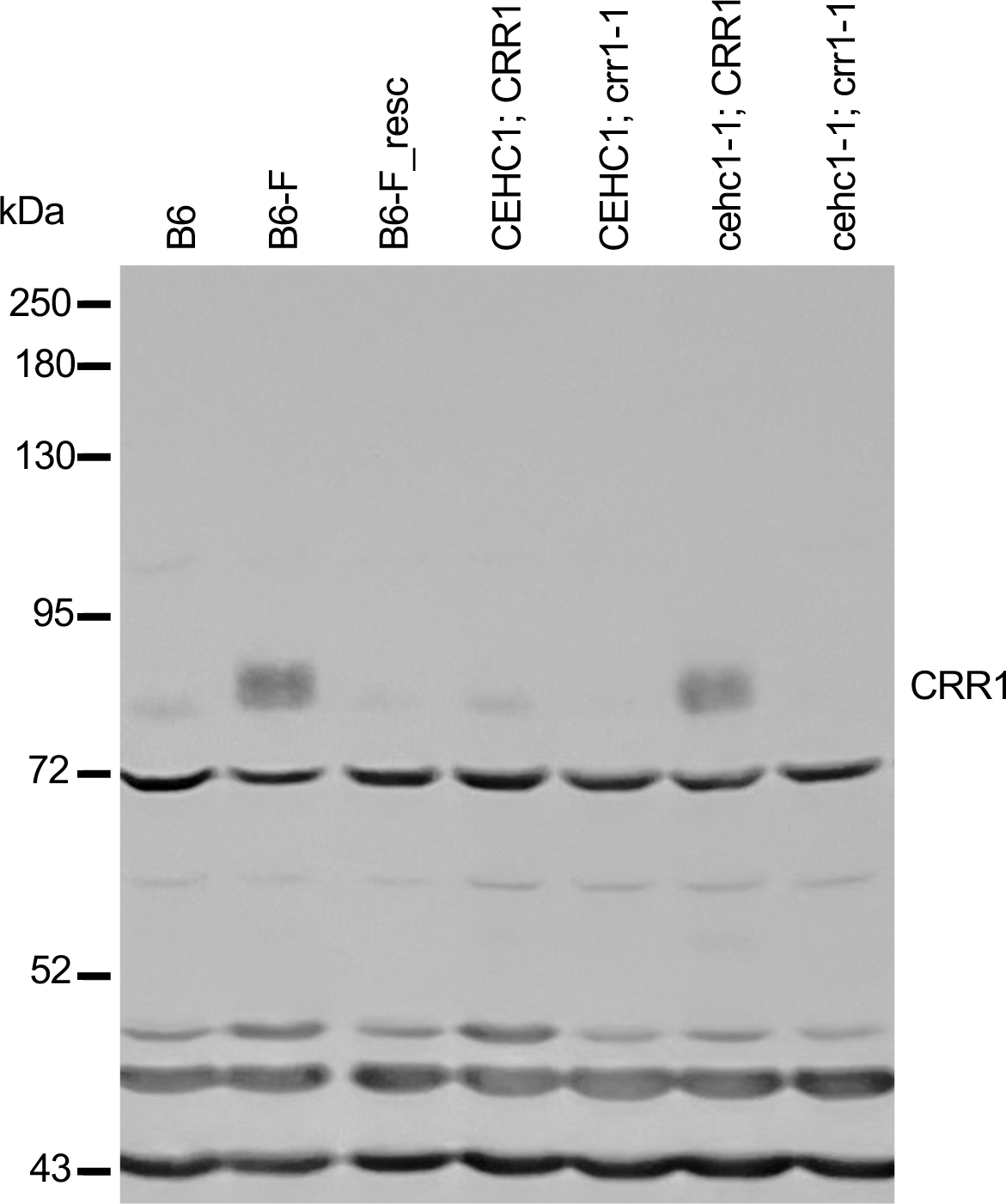
Full immunoblot of Figure 8. Image of the full immunoblot used in Figure 8C. The immunoblot was prepared using Protein Gels and Immunoblotting Method 1 with CRR1-B antiserum.

**Supplementary Figure 5.**
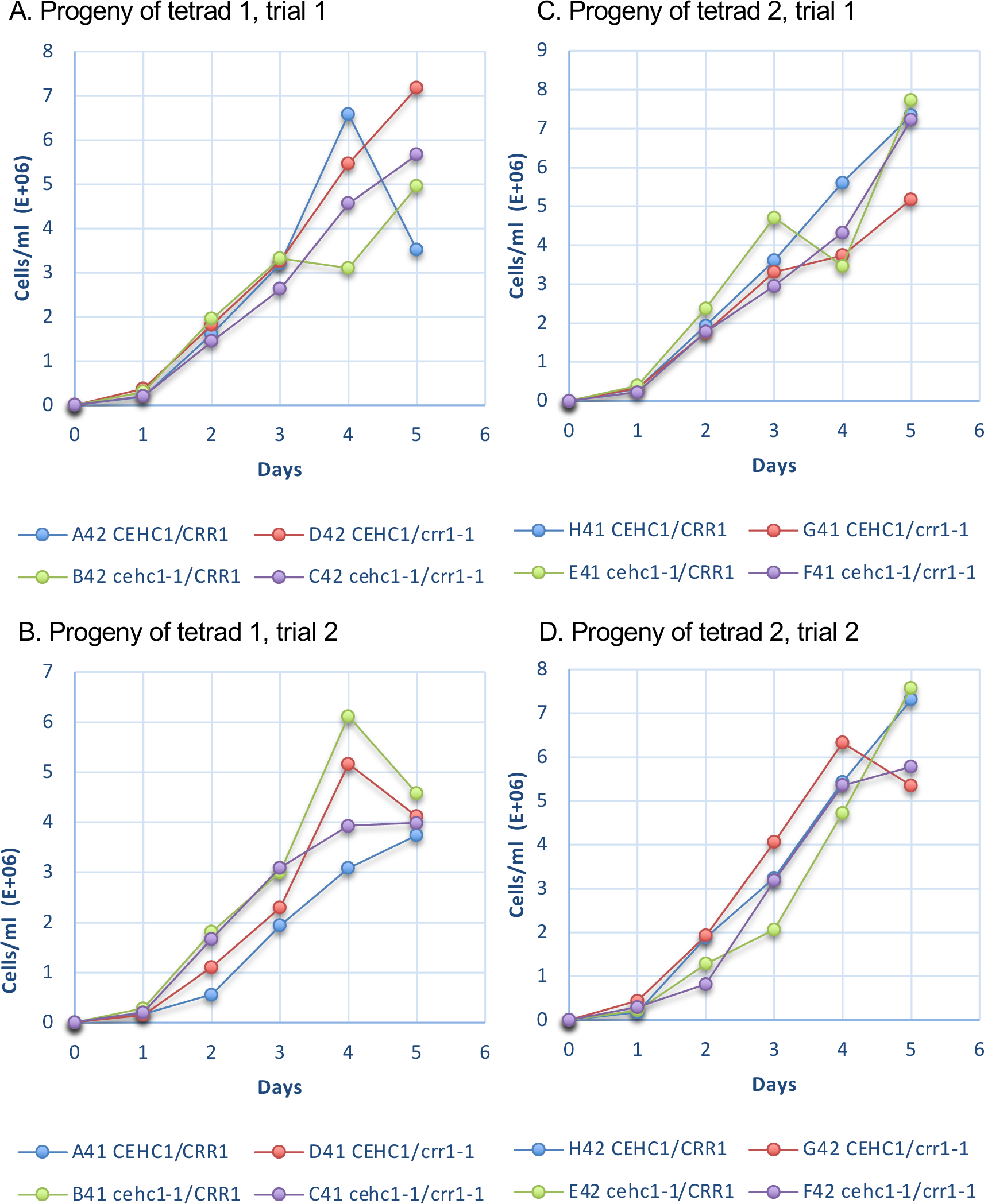
Effect of *CHC1*, *CRR1* mutant and wild-type alleles on cell growth. Two complete tetrads from a cross of CC-125 x CS1216C6 (Supplementary Table 1) were genotyped (Supplementary Table 5). Cells were grown in liquid minimal medium and inoculated into 20 ml cultures in 250 ml flasks at an initial density of 2 – 6 × 10^3^ (day 0). Cultures were rotated on an orbit shaker for aeration and illuminated with 10,000 lx with constant lighting from cool white fluorescent bulbs. Samples were collected at 24h intervals and cells were fixed for counting with a hemocytometer. Two independent experiments were carried out with each set of tetrads.

